# Site-specific Effector Protein Functionalization to Create Bead-based Avidity Model Systems

**DOI:** 10.1101/2023.03.27.534459

**Authors:** Markus Hackl, Dharanidaran Jayachandran, Khovesh Ramdin, Tong Zhong, Timothy Travers, Sandrasegaram Gnanakaran, Shishir P.S. Chundawat

**Author notes:** **Corresponding author:** Shishir P. S. Chundawat. Department of Chemical & Biochemical Engineering, Rutgers, The State University of New Jersey, 98 Brett Road, Piscataway, NJ 08854. **Email:**.

## Abstract

The cooperative effect of multiple affinity binding interactions creating a stable bond, known as avidity, is a universal biological phenomenon seen in diverse systems. For example, avidity based biomolecular interactions are particularly important in assessing the potency of potential drugs such as monoclonal antibodies, chimeric antigen receptor (CAR) T-cell, or Natural Killer, cells to treat cancer or engineering microbes with cell surface immobilized enzyme complexes for consolidated bioprocessing (CBP) of cellulosic biomass to fuels and chemicals. However, predicting or measuring avidity based on *in vitro* single affinity interactions with non-complexed protein-ligand binding model systems has limitations and often fails to describe the avidity effects observed *in vivo* with cell surface complexed proteins interacting with multivalent ligands at solid interfaces. Acoustic force spectroscopy (AFS) based assays have recently emerged as a reliable method for direct avidity measurements, expressed as adhesion or rupture forces, which positively correlate with *in vivo* avidity interactions. However, to better understand and model avidity, in particular for cell-cell interactions and to correlate it with classical binding affinity, a cell mimetic model system with controlled avidity-related properties is needed. Here, we present a method for producing such a cell mimetic model system using “effector beads” that can be used in AFS-based avidity assays or any other bead-based avidity assay. The protein of interest is heterologously expressed and biotinylated *in vivo* in *E. coli*, purified, and subsequently tethered with streptavidin coated micron-sized beads to create effector beads. Our experimental results, combined with simulations of the multivalent binding phenomena, demonstrate the dependency of bead rupture force on its receptor protein surface density and force loading rate as well as the intrinsic kinetic binding parameters of the protein-ligand system of interest. These insights provide valuable information for designing future effector bead assays and cell avidity measurements for screening and characterization purposes for diverse applications.

## Introduction

Multivalent protein-ligand binding interactions are important in many biological systems. In biofuel production, the addition of protein domains such as carbohydrate binding modules (CBM) to catalytic domains of enzymes or the assembly of larger CBM/catalytic domain complexes on microbial cell surfaces called cellulosomes increases the synergistic efficiency for enzymatic deconstruction of cellulosic biomass into fermentable sugars (1–5). Another example is the creation of strong non-covalent bonds between immunogenic host cells or exogenous pathogen cells with the endothelial cell lining tissues in the presence of flow induced shear stress due to the accumulation of many low-affinity binding cell-surface interactions (6–8). These microscopic binding events translating to macroscopic binding phenomena are often termed avidity or functional avidity (9).

Avidity effects are particularly important in assessing the potency of potential drugs such as monoclonal antibodies (10) as well as chimeric antigen receptor T (CAR T) cells and natural killer (NK) cells to treat various cancers (11–13). Due to synergistic multivalency and cooperative effects, the measured avidity is often higher than the sum of the individual binding affinities of its constituents (9, 14–16). This makes the prediction of avidity based on single affinity measurements difficult if all molecular level interactions are not identified a priori. Several methods have been developed to assess avidity interactions such as atomic force microscopy (17), surface plasmon resonance (18, 19), or shear force induced methods (20– 22). However, these methods either lack high throughput, are time-consuming, or difficult to reproduce.

Acoustic force-based avidity assays have recently emerged as an alternative method to reliably assess avidity effects of cells, in particular for CAR T cell engineering and characterization (23–29). Such avidity assays are simple to execute and provide reliable predictions for T-cell efficacy. However, the underlying single affinity interactions are not well characterized since it is difficult to control or determine the exact number of interactions between cells and its surface targets.

One way to better understand avidity for cell-cell or cell-surface interactions and correlate it with affinity is to create a cell mimetic model system using functionalized beads (or effector beads), which display controlled avidity related properties such as receptor density and receptor orientation. Effector-functionalized beads are used, for example, in flow cytometry (30, 31) or in expanding T-cells (32, 33). However, the commercial availability of either effector-functionalized beads or biotinylated antibodies to create effector beads is limited. In this study, we describe a universal method for creating site-specifically biotinylated effector proteins for effector bead functionalization and subsequent use in acoustic force spectroscopy-based avidity assays. As a proof of concept, we used streptavidin-coated polystyrene particles that were functionalized with biotinylated carbohydrate binding modules (CBM) as the model receptor and a cellulose surface as the model ligand. We inserted the AviTag (34) on the N terminus of the proteins and used an *in vivo* biotinylation approach in *E. coli* to produce high purity biotinylated proteins in a single step. The assay execution is simple and data analysis is straightforward with a customized, open-source MATLAB-based graphical user interface (GUI) developed for users with limited experience with force spectroscopy techniques. A visual representation of the overall assay workflow is shown in **Figure 1**. Our results show that the relative avidity (or rupture force) is dependent on the surface density of receptors on the beads as well as the force loading rate, which provides valuable information for designing future effector bead assays and screening antigen targets in the early stages of drug discovery or for development of more efficient cellulolytic enzymes or microbes for biofuels production.

**Figure 1:**
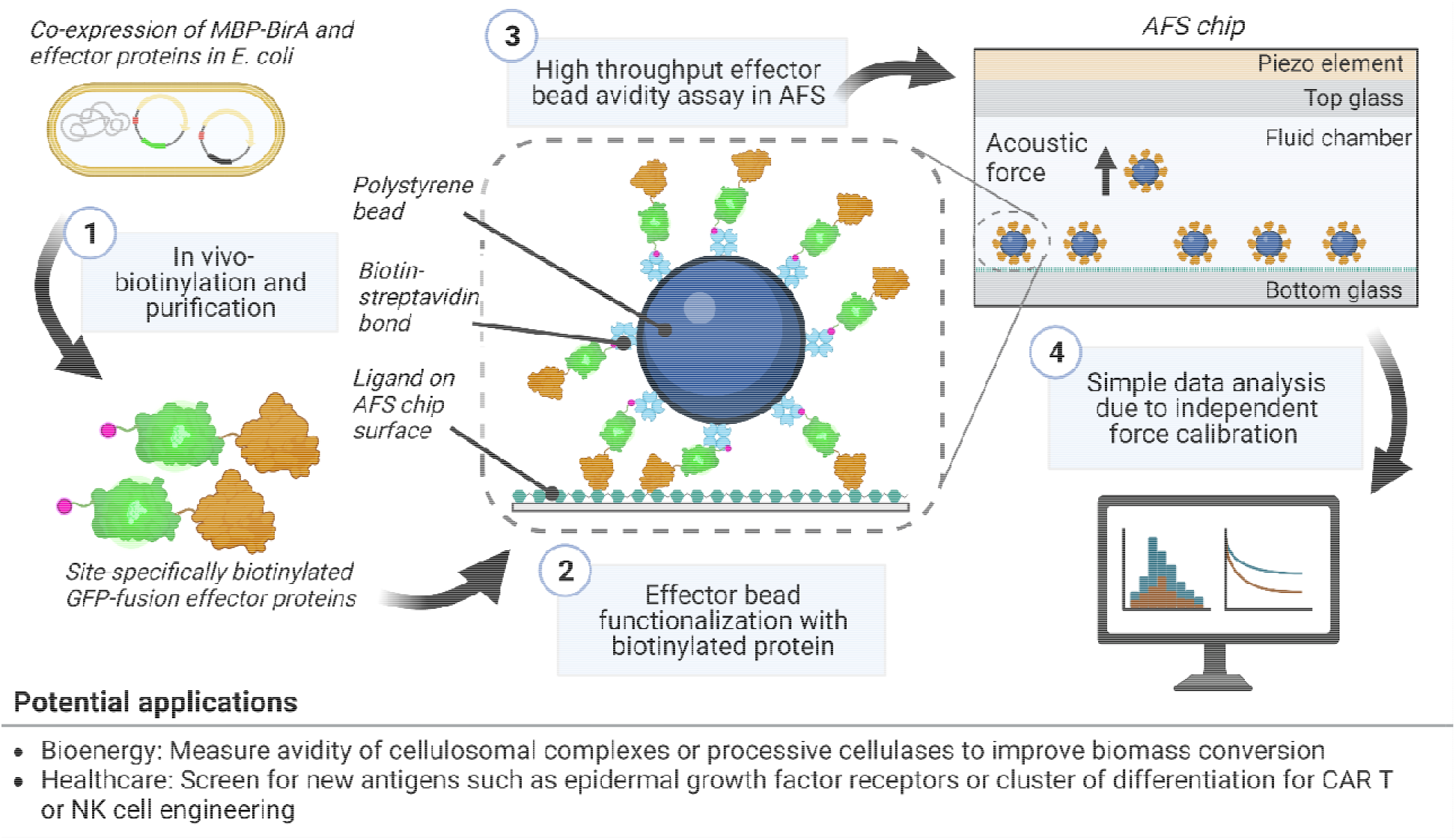
Overview of workflow to create effector beads for avidity assays using AFS. Step 1: The plasmid containing an Avi-tagged green fluorescent protein (GFP)-fused effector protein is co-transformed with a plasmid for Biotin ligase (BirA) fused to maltose binding protein (MBP) in E. coli. Co-expression of BirA results in most Avi-tagged fusion proteins being biotinylated before purification. Step 2: Streptavidin-coated polystyrene beads are then functionalized with effector proteins and allowed to bind to a ligand-functionalized AFS chip surface. Step 3: Acoustic forces are applied at varying loading rates which pull the bound effector beads away from the surface, thus creating force-induced unbinding events. Step 4: The data can be quickly analyzed with our developed graphical user interface (GUI).

## Materials and Methods

### Reagents & Buffers

#### Chemicals and substrates

Unless otherwise mentioned, all reagents were either purchased from VWR International, Fisher Scientific, or Sigma-Aldrich. Streptavidin-coated beads (nominal diameter 5.14 μm) were purchased from Spherotech, Inc. Sulfuric acid-hydrolyzed nanocrystalline cellulose (NCC) was kindly donated b Richard Reiner from the USDA Forest Products Laboratory (Madison, Wisconsin) (35).

#### Buffers

Two buffers were used to block the AFS chip surface before any experiments. Buffer B1 consists of 10 mM phosphate buffer (PB) at pH 7.4 supplemented with 2.5 mg/ml bovine serum albumin (BSA) and casein respectively. Buffer B2 consists of 10 mM PB supplemented with 2.5 mg/ml BSA and casein, respectively, and 5.6 mg/ml Pluronic F-127 as previously described (36). During the bead functionalization and washing steps, a working buffer (WB) was used consisting of 10 mM PB supplemented with 0.31 mg/ml BSA and casein and 0.19 mg/ml Pluronic F-127, respectively. All rupture force experiments were carried out in 10 mM PB without any additional supplements. All buffers were degassed at -90kPa for 15-30 minutes before use each time. In addition, Pro-Clin 300 at a concentration of 0.05% v/v was added to all buffers to suppress potential growth of microbes, since the same AFS NCC chip can be typically reused over 2 weeks.

### In vivo biotinylation of carbohydrate binding modules

#### Vector modification

The plasmids harboring *Clostridium thermocellum* CBM3a and *Spirochaeta thermophila* CBM64 carbohydrate-binding modules fused to green fluorescent protein (GFP) (herein referred to as CBM3a and CBM64 respectively) were prepared as previously described (37). The tandem versions of both proteins (GFP-CBM-CBM) were prepared as previously described (38). The additional CBM domain is added to the C terminus using the same flexible linker as between the GFP domain and first CBM motif to create the tandem CBM constructs.

The histidine tag for purification is located at the N-terminus of the sequence, followed by a Tobacco Etch Virus (TEV) cleavage site, then GFP with a linker domain and CBM. The AviTag was inserted between the TEV cleavage site and GFP by amplifying the vector using primers containing the AviTag sequence with a 15 bp overhang complementary to the vector backbone. The amplicon was transformed into *E. cloni* 10G cells using sequence and ligation independent cloning (SLIC) (39). The insertion of the AviTag was confirmed by sequencing the plasmid extracted from *E. cloni*. The sequence of tandem GFP-CBM3a is shown in Supplemental Table S1.

The vector encoding BirA fused to a maltose binding protein (MBP), pH6-MBP-TEV-BirA, was a gift from Mark Arbing (Addgene plasmid # 179694). The histidine tag of the MBP-BirA plasmid was deleted using SLIC (39), since the CBMs needed to be purified using the same tag. The primer sequences used to insert the AviTag into the GFP-CBM constructs and remove the histidine tag from MBP-BirA are listed in Supplemental Table S2.

#### Protein expression and purification

The CBM plasmids were transformed into chemically competent *E. coli* BL21-CodonPlus-RIPL [DE3] already containing the MBP-BirA plasmid and grown at 37°C as a 10 ml overnight culture in LB media containing 34 μg/ml chloramphenicol, 100 μg/ml carbenicillin and 50 μg/ml kanamycin. From this overnight culture, 2 ml starter volume was used to inoculate 50 ml LB media containing the same antibiotics and cells were grown at 37°C with shaking at 200 rpm. Once the optical density (OD_600_) reached 0.4-0.6, the expression was induced by adding 1 mM isopropyl ß-D-1-thiogalactopyranoside (IPTG) and 200 μM biotin, and reducing the temperature to 25°C. After approximately 18 hours of incubation, the cells were harvested by centrifugation at 6,000 x g for 15 minutes.

The cell pellets (∼0.5-1 g) were resuspended in 7 ml lysis buffer (10mM PB at pH 7.4 containing 100mM sodium chloride, 20% v/v glycerol and 0.3% v/v Triton X-100) and lysed by sonication on ice. Next, the cells were separated from the supernatant containing the biotinylated GFP-CBM constructs by centrifugation at 10,000 x g for 40 minutes at 4°C. The supernatant was mixed with 1 ml of nickel-nitrilotriacetic acid (Ni-NTA) functionalized magnetic beads and incubated for 60 minutes at 4°C with gentle mixing. Next, the supernatant was removed, and the resin was incubated with 20 ml lysis buffer for 20-30 minutes followed by incubation in 20 ml of 10 mM phosphate buffer (supplemented with 0.1% v/v Triton X-100) for 20-30 minutes. The resin was washed twice with 10 ml IMAC-A buffer (100 mM 3- (N-morpholino) propanesulfonic acid (MOPS), 10 mM imidazole, 500 mM sodium chloride and 0.1 % v/v Triton X-100 at pH 7.4) and twice with 10 ml buffer containing 95% IMAC-A and 5% IMAC-B (100 mM MOPS, 500 mM imidazole and 500 mM sodium chloride at pH 7.4). Finally, the GFP-CBM constructs were eluted from the resin by incubation in 1 ml IMAC-B for 10-15 minutes at 4°C. The purified protein concentrations were determined by measuring absorbances at 280 nm using the molar extinction coefficients listed in Supplemental Table S3.

#### Extent of protein biotinylation using gel-shift assay

The extend of biotinylation of the Avi-tagged CBMs was confirmed using a gel-shift assay (40). Briefly, the purified proteins were diluted to 3 μM in 10 mM PB, pH 7.4. In two PCR tubes for each CBM construct, 5 μl of protein were mixed with 10 μl of 2x Laemmli buffer (lacking ß-mercaptoethanol) and incubated at 95°C for 5 minutes. After the samples cooled to room temperature, one PCR tube was mixed with 5 μl of PB whereas the other sample was mixed with 5 μl of 3 μM streptavidin (SA) dissolved in PB. The samples were incubated at room temperature for an additional 5-10 minutes. Next, 15 μl of each sample was run on a 10% polyacrylamide gel and stained with Coomassie Blue. An additional sample for each protein was prepared using 2x Laemmli containing ß-mercaptoethanol to qualitatively assess protein purity.

The extent of biotinylation was estimated using densitometry by comparing the intensity of the streptavidin-containing band to the band of the non-streptavidin sample using GelDoc Image Lab Software (BioRad, Hercules, CA). Considering the extent of biotinylation, the GFP-CBM constructs were diluted to 10 μM biotinylated protein, aliquoted and stored at -80°C.

### Development of easy-to-use graphical user interface for AFS force calibration and rupture data analysis

#### Force calibration overview

Typically, beads are tethered to the AFS chip surface using a DNA tether and a force calibration is performed on each bead before executing a rupture force protocol (41). However, the avidity assays described here do not contain DNA tethers and an alternative force calibration needed to be performed to correlate the applied amplitude (power) with the force experienced by each bead. This force calibration step is necessary because the force experienced by the bead varies with the bead’s position within the AFS chip (42). Although a software for the tether-free stokes force calibration (SFC) was previously developed (42), it was not compatible with the updated tracking software and file format provided by LUMICKS B.V. The updated tracking software uses a power value (expressed as a value between 0-100%) as the control parameter instead of the amplitude as previously described (41, 43) and the power value corresponds linearly to the applied force. Hence, the force calibration implemented in a MATLAB-based graphical user interface (GUI) presented here follows equation 1 to relate the applied power (*P*) to the location-dependent acoustic force (*F*_*ac*_) exerted on particles inside the AFS chip.

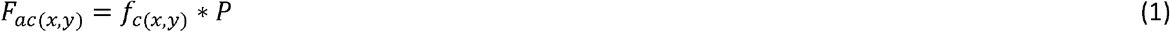

Where *f*_*c*_ represents the force calibration factor for a particular location within the field of view (FoV). The approximated force equilibrium of a bead subjected to acoustic forces, rearranged to express the bead velocity (42) is given in equation 2 as,

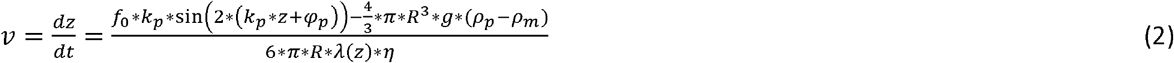

Similar to the previous analysis software (42), equation 2 was numerically integrated using the analytical Brenner correction for the viscosity (*λ* (*z*) \**η*) to optimize the force fit parameters *f*_*0*_, *k*_*p*_ and *φ* _*p*_ for a given bead radius, *R*, and particle and media density *ρ* _*p*_ and *ρ* _*m*_, respectively. A detailed description of all software features as well as detailed steps for data analysis is available in the Supplemental Information.

#### Creation of force factor heat map for each functionalized AFS chip

The force factor heat map represents the location dependence of the force calibration factor (**Figure 3**-C). All experiments were performed on a G1 AFS instrument with G2 AFS chips provided by LUMICKS B.V. The SFC was performed after immobilization of nanocellulose within the chip (as described below). First, the chip was rinsed with 0.5 ml deionized (DI) water followed by 0.5 ml of PB. To reduce non-specific binding of the streptavidin-coated beads, the surface was passivated in B1 and B2 for 15 minutes each. Next, the chip was rinsed with 0.2-0.5 ml of PB. The beads (5.14 μm diameter) were diluted in WB at a ratio of approximately 1:50 (v/v) to yield between 80-160 beads in the FoV at 10x magnification. An increase in the bead concentration for SFC tends to reduce the fraction of useful traces since the beads were slowly pushed towards the closest pressure node (42) where their overlap in diffraction patterns interfered with the accurate 3D position determination.

The beads were allowed to settle for 5 minutes. Next, a look-up-table (LUT) was created to determine their z-position perpendicular to the chip surface (41). During SFC, beads were tracked with a moving region of interest (ROI) so that multiple SFC runs could be performed with the same set of beads. Beads were tracked at a frame rate of 35-40 Hz using a 10x magnification objective. During the first 20-30 seconds, the diffusion of the beads was recorded in the absence of force to determine the anchor point (41). Next, a constant, small force (0.5-1.5 pN) was applied for 1 second. The force applied was high enough to lift the bead up to the acoustic pressure node (z-node), but small enough so that the tracking at the relatively low frame rate produced enough data points at the time frame where the bead was rising. If those conditions were met, no iterative force fit and z-node estimation was needed as previously described (42). Typically, the SFC was performed with 3 distinct power values, with at least 1000 beads recorded for each power value (between 40-60 individual SFC experiments).

The developed GUI automatically reads in the tracking data, performs the SFC fit and removes outliers based on the following criteria: (i) tracking errors as defined by standard deviation in z of more than 2 μm during anchor point determination, (ii) z-position of < 10 μm during force application indicating weak force or stuck bead, (iii) R^2^ value during SFC fit of less than 0.99, (iv) Z-score > 3 of z-node or fit parameters, (v) z-node position outside 15-25 μm. Beads were merged based for a segment size of 10 × 10 μm (42) for the FoV (679 × 543 μm at 10x magnification). Segments in which a higher power value yielded a lower force compared to lower power values, were removed as well (42). Equation 1 was fitted to the data to obtain the force calibration factor, *f*_*c*_, if at least two beads were present in a particular segment. Once the heatmap was established, it was used for all experiments in the same FoV of a given AFS chip.

### Characterizing effector bead avidity

#### Influence of effector concentration on rupture force and initial binding commitment

The NCC ligand film was deposited on the AFS chip surface using an automated protocol described previously (36). To functionalize beads with CBMs, 10 μl of CBM3a diluted in WB at various concentrations were added to 5 μl of streptavidin coated beads and mixed on a rotisserie shaker for 15 minutes. The beads were washed twice with 100 μl WB to remove any unbound proteins, diluted in 20-40 μl PB and inserted into the surface-passivated AFS chip. Beads were allowed to bind to the surface for 15 minutes after which unbound beads were removed by flushing the chip with PB at a flow rate of 3 μl/min and a constant applied acoustic force of 0.1-0.2 pN.

Similar to the SFC protocol above, a LUT was created and beads were tracked, however with a fixed ROI and a frame rate of 20 Hz. For the first 60 seconds, the beads were tracked in the absence of force to determine the anchor point. Next, a linear force ramp of 1 pN/s (based on the mean *f*_*c*_ value determined by SFC) was applied until all beads ruptured or 100% power was reached. Each experiment was carried out in at least duplicate and until N>50 was reached for each CBM concentration.

Bead traces were analyzed using the customized GUI software developed in this work. After removing outliers based on a standard deviation of > 2 μm in z-direction during the anchor point determination, each z-trace of the remaining beads was averaged using a moving window of 50-150 frames. The GUI automatically determines the rupture point based on a user-defined z-value cut-off. However, some beads may show tracking errors during the force ramp due to beads moving over a tracked ROI. Such beads were flagged by the GUI and their rupture time was determined manually. Based on the heatmap created above, the rupture force was determined by obtaining an interpolated force calibration factor for the respective bead position, which was multiplied by the power value at bead rupture time according to equation 1. The detailed description for rupture force determination is found in the Supplemental Information.

#### Loading rate dependence and binding percentage of effector-functionalized beads

Beads were functionalized by incubating 10 μl of 500 nM CBM (CBM3a, tandem CBM3a, CBM64 and tandem CBM64 respectively) diluted in WB with 5 μl of streptavidin coated beads for 15 minutes. The washing steps, bead incubation and tracking were carried out as mentioned above. Experiments below a loading rate of 10 pN/s were recorded at a frame rate between 10 and 20 Hz whereas 40 Hz was used for a loading rate of 100 pN/s.

The loading rate was varied between 0.01-100 pN/s and rupture force experiments were carried out in triplicate for each CBM construct and loading rate. The analysis of traces was carried out as described above. Rupture force data for each loading rate was pooled together for the same loading rate and the mean loading rate was determined by fitting a normal distribution to the loading rate histogram. However, similar to our previous CBM-cellulose rupture force assay for single molecules (36), all 4 CBMs showed a tail towards larger rupture forces. Hence, a lognormal distribution was fitted to the rupture force data to obtain a representative mean value for each protein construct. For the generation of the bound beads as a function of the applied force, the data was not pooled, but the mean and average deviation of 3 experiments was determined.

For re-use and short-term storage at room temperature, the NCC ligand functionalized chips were rinsed with 0.2 ml of 0.5 % (v/v) bleach, followed by 0.5 ml of 0.5 M sodium thiosulfate and 2 ml of DI water. The microfluidic chamber was filled with DI water supplemented with 0.05% v/v Pro-Clin 300 and sealed with parafilm to prevent drying.

### Modeling the multivalent unbinding of CBM-functionalized beads from cellulose surface

The effective equilibrium association constant for binding of two domains from the same molecule to another molecule containing multiple ligand sites can be given by:

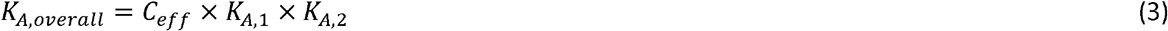

Where, *K*_*A,1*_, and *K*_*A,2*_, are the equilibrium association constants of the individual domains and *C*_*eff*_ represents the effective concentration of unbound ligand experienced by the unbound domain (after binding of the other domain). To account for the cellulose polymers being restricted to a surface instead of being free-floating, we incorporated a volume-based corrective factor (for detailed description, see Supplemental Information) that reduces the *C*_*eff*_ (44, 45) of cellulose experienced by the unbound CBM in a tandem construct upon binding of the other CBM:

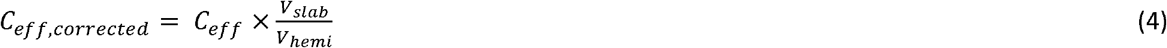

with *V*_*slab*_ = 4 *π r*^*2*^*h* and 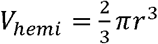 where *r* = 2.5 *μm* represents the radius of the polystyrene bead to which CBM constructs are functionalized, and *h* = 6 *nm* the average diameter of a cellulose microfibril (46). The binding enhancement factors calculated above were used for developing a simplified systems model for beads that have been functionalized with either single or tandem CBM constructs under varying force loading rate and protein construct density on the bead surface. All simulations of this model begin with an equilibration where CBM-functionalized beads can reversibly bind to cellulose polymers (see Supplemental Information). The steady-state levels obtained after equilibration are then used as the starting point for the subsequent loading rate application phase where the rupture force is increased over time from 0 to 200 pN. Beads that detach from cellulose during loading rate application in the simulations are not allowed to rebind as detached beads in the AFS experiments also stay unbound due to the applied acoustic force, which keeps beads away from the surface.

Because of the combinatorial complexity of the system in terms of the number of potential interactions, it is challenging to specify all possible reactions in the protein-ligand binding network. We therefore performed the modeling using BioNetGen (47, 48) that is capable of generating the entire reaction network for a system from a smaller number of reaction rules (see Supplemental Information). Each rule specifies a pattern or set of patterns that are shared by multiple reactions in the network. For instance, one rule can specify all reversible binding events for the first CBM moiety from the bead to cellulose during the equilibration phase (equation S11, Supplemental Information) which involve bimolecular complex formation. Whereas another rule can specify all reversible tandem binding effects of the second CBM moiety in a bead-cellulose complex (equation S12, Supplemental Information). Note that the binding reactions generated from the latter rule are modeled with unimolecular rates since the CBM-functionalized bead is already in complex with cellulose due to prior binding of at least one CBM moiety from the bead.

## Results and Discussion

### In vivo biotinylation of carbohydrate binding modules

Fusion of MBP to BirA resulted in reproducible *in vivo* biotinylation of heterologously expressed proteins in *E. coli* (49). Previous reports indicated a reduction in the growth rate of *E. coli* when BirA is overexpressed (50), however under conditions described here, no significant reduction in the growth rate was observed (Supplemental Figure S1). Using our previously described purification conditions (51) resulted in a stable complex of MBP-BirA and GFP-CBMs during IMAC purification (Supplemental Figure S2, lane 2), most likely due to the presence of salts (500 mM) and absence of polymers which are known to reduce protein-protein interactions. Thus, the concentration of sodium chloride in the lysis and washing buffers was reduced to 100 mM. Furthermore, Triton X-100 was added to both buffers, which was shown to reduce the interaction between BirA and the AviTag (Supplemental Figure S2, Lane 13). From a 50 ml cell culture, we obtained between 2-4 mg (> 35 nmol) of IMAC purified, biotinylated proteins.

With the described purification method, all target proteins are separated from the cell lysate without the detection of any by-products as shown in **Figure 2**, lane 11-14. CBM3a (lane 2) and tandem CBM3a (lane 6) show faint bands at higher and lower molecular weights, most likely due to incomplete unfolding under non-reducing conditions, since only one band is observed under reducing conditions (lanes 11 and 13, respectively). Streptavidin-containing lanes 3, 5, 7, 9 for CBM3a, CBM64, tandem CBM3a and tandem CBM64 respectively, show higher molecular-weight species due to the complex formation of biotinylated CBMs with streptavidin. The extent of biotinylation determined by densitometric analysis was > 95% for CBM3a, CBM64, and tandem CBM3a and ∼80% for tandem CBM64, respectively. Although biotin was provided in excess (200 μM in growth medium during induction with IPTG), a 100% biotinylation rate was not achieved. Since the protein concentration was adjusted to the amount of biotinylated products, incomplete *in vivo* biotinylation has limited effect on any subsequent bead functionalization steps. If only the presence of biotinylated products is required, the IMAC-purified proteins can be further purified using monomeric avidin-functionalized resin, which exhibit a lower affinity for biotin compared to tetrameric streptavidin (52, 53).

**Figure 2:**
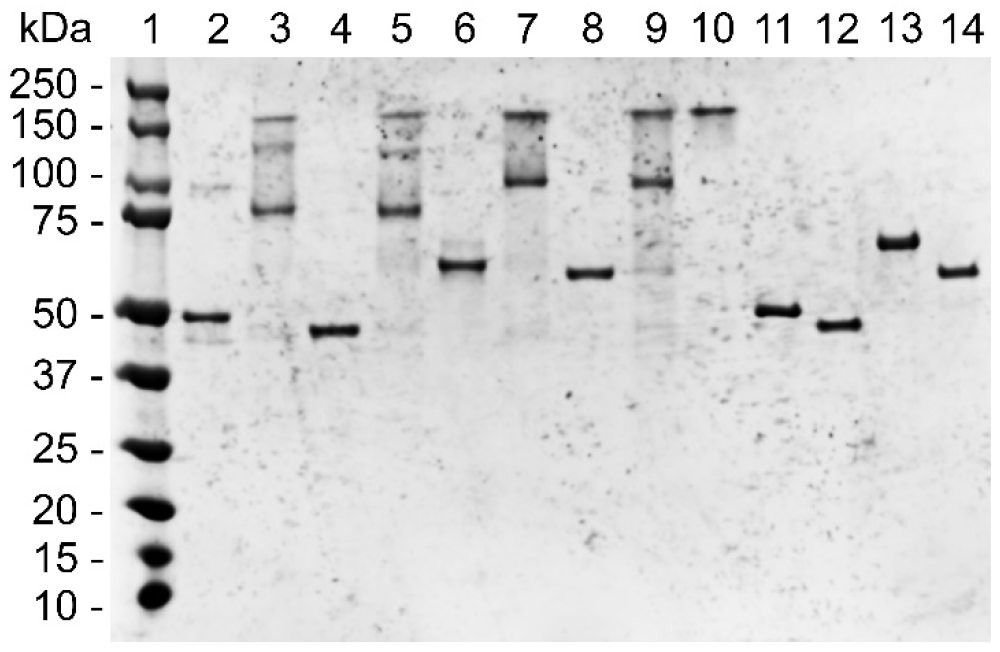
SDS-PAGE gel image shows the extent of biotinylation and protein purity. Samples for lanes 2-10 were incubated under non-reducing conditions, lanes 11-14 under reducing conditions. SA stands for Streptavidin. Lane 1: protein standard; Lane 2: GFP-CBM3a (no SA); Lane 3: GFP-CBM3a (with SA); Lane 4: GFP-CBM64 (no SA); Lane 5: GFP-CBM64 (with SA); Lane 6: tandem GFP-CBM3a (no SA); Lane 7: tandem GFP-CBM3a (with SA); Lane 8: tandem GFP-CBM64 (no SA; Lane 9: tandem GFP-CBM64 (with SA); Lane 10: SA only; Lane 11: GFP-CBM3a; Lane 12: GFP-CBM64; Lane 13: tandem GFP-CBM3a; Lane 14: tandem GFP-CBM64.

### Development of easy-to-use graphical user interface for AFS force calibration and rupture data analysis

**Figure 3**-A shows the z data of 10 randomly selected beads during the time range considered for SFC, when power was turned on and beads were lifted in z towards the z-node. The blue lines in **Figure 3**-A indicates the fit of the numeric integration of equation 2. All fits exhibit R^2^ > 0.99. The dependence of force on the z-position for those 10 traces is visualized in **Figure 3**-B, indicating that the z-node is approximately 20 μm above the surface. **Figure 3**-C displays the force factor heatmap where 700 merged ROIs were obtained (black circles) and the location of the 10 randomly selected traces (red dots) shown in **Figure 3**-A and B. As previously characterized (42), the exerted force varies greatly within the FoV, highlighting the importance of obtaining a location dependent force calibration.

**Figure 3:**
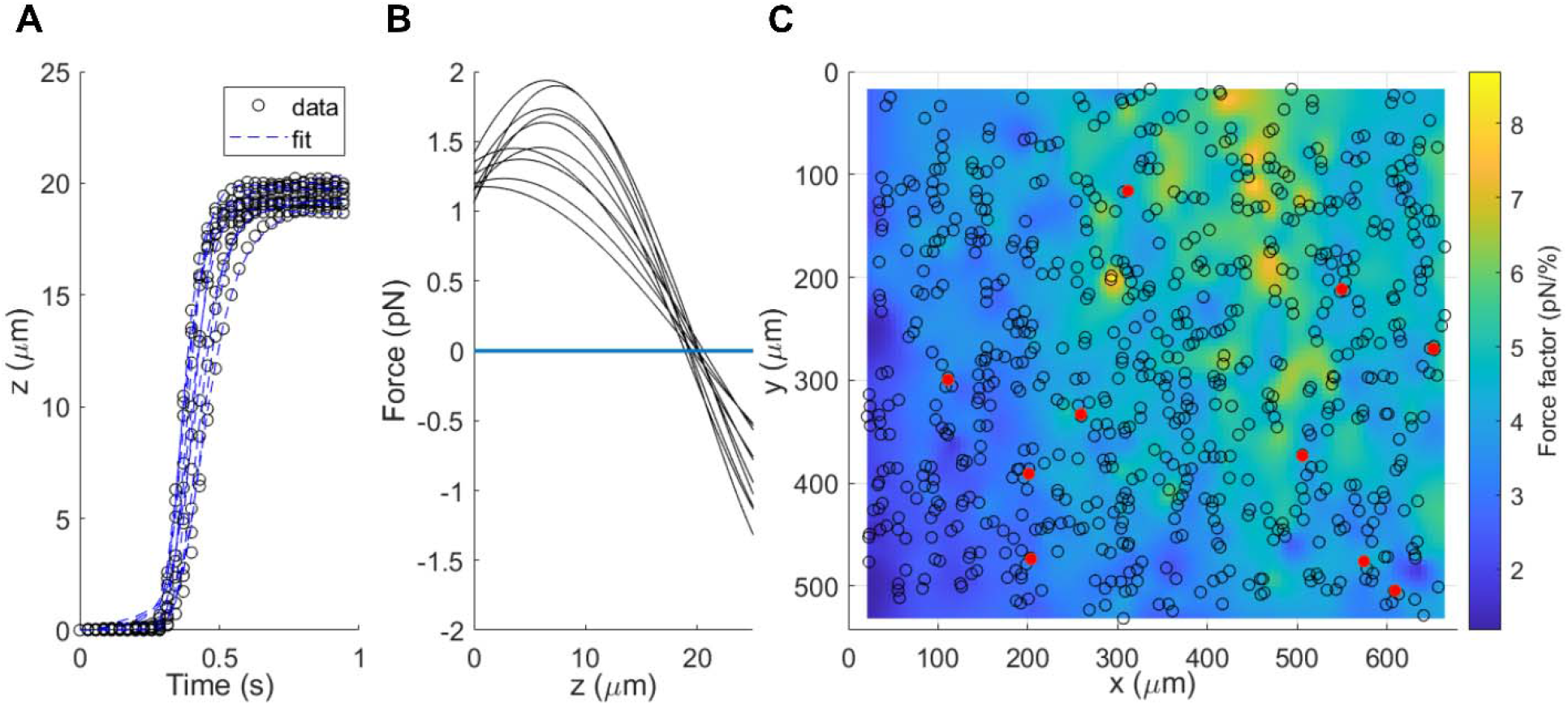
Developed GUI enables quick and in-depth analysis of bead traces during stokes force calibration (SFC). A) z trace of 10 randomly selected beads during SFC (power applied so that beads are lifted towards the z-node). The black circles show the raw data, and the dashed blue lines show the fit of the numerical integration of equation 2. B) Force profile of the 10 traces visualizing the change in force as a function of the distance of the bead from the surface as well as the position of the z-node. The guide at F=0 is shown as a visual aid. C) Force factor heatmap NCC-functionalized AFS chip showing the variation of effective force over the FoV. The merged ROIs are shown as black circles, whereas the 10 randomly selected traces from A) and B) are shown as red dots. The mean force calibration factor is 4.1 ± 1.0 pN/% (μ ± σ of normal fit), highlighting the importance of location-specific force calibration.

The user-friendly GUI developed here allows for efficient analysis of bead traces with the click of a few buttons. For example, during SFC, the fit of individual traces as well as the force dependence in z can be readily visualized (**Figure 3**-A and B, Supplemental Figure S3). Furthermore, the rupture forces are determined automatically, and only a fraction of traces (< 10%) need to be analyzed manually. The GUI automatically assigns the rupture force based on a pre-determined force factor heatmap and visualizes the data in histograms (Supplemental Figure S4). A thorough description of the GUI features along with the source code and training data is found at github.com/ChundawatLab/AFS-GUI (54).

### Characterizing effector bead avidity using force spectroscopy

#### Influence of effector concentration on rupture force and initial binding commitment

Because the beads have a limited number of available biotin binding sites on the beads are limited (0.8 μM or 1.2*10^−6^ pmol/bead), we aimed to investigate how the sensitivity of the rupture force varies with respect to the CBM concentration used to functionalize beads. **Figure 4**-A shows the influence of the used CBM concentration on the rupture force. With the current protocol (10 μl of CBM dilution mixed with 5 μl of beads), a 100nM CBM dilution corresponds to a molar excess of 0.25 with respect to available biotin binding sites on the beads. The equimolar concentration is reached at 400 nM of CBM dilution. Above a concentration of 100 nM, the rupture force remains unchanged, but drops significantly for CBM concentrations below 100 nM as seen in **Figure 4**-A. The mean rupture forces for <10 nM are within the range of previous single-molecule experiments (36), however, a direct comparison is difficult to achieve, since the exact number of CBMs interacting with the surface at once is not known. The ratio of the number of beads tracked vs the number of beads initially in the FoV before the flushing step is expressed as the initial binding commitment in **Figure 4**-A. Using streptavidin-coated beads only as a control, the initial binding commitment was <1% and the average rupture force <5 pN. The initial binding commitment remains >90% for CBM concentrations greater than 15 nM, but then drops below 50% for 10 nM.

**Figure 4:**
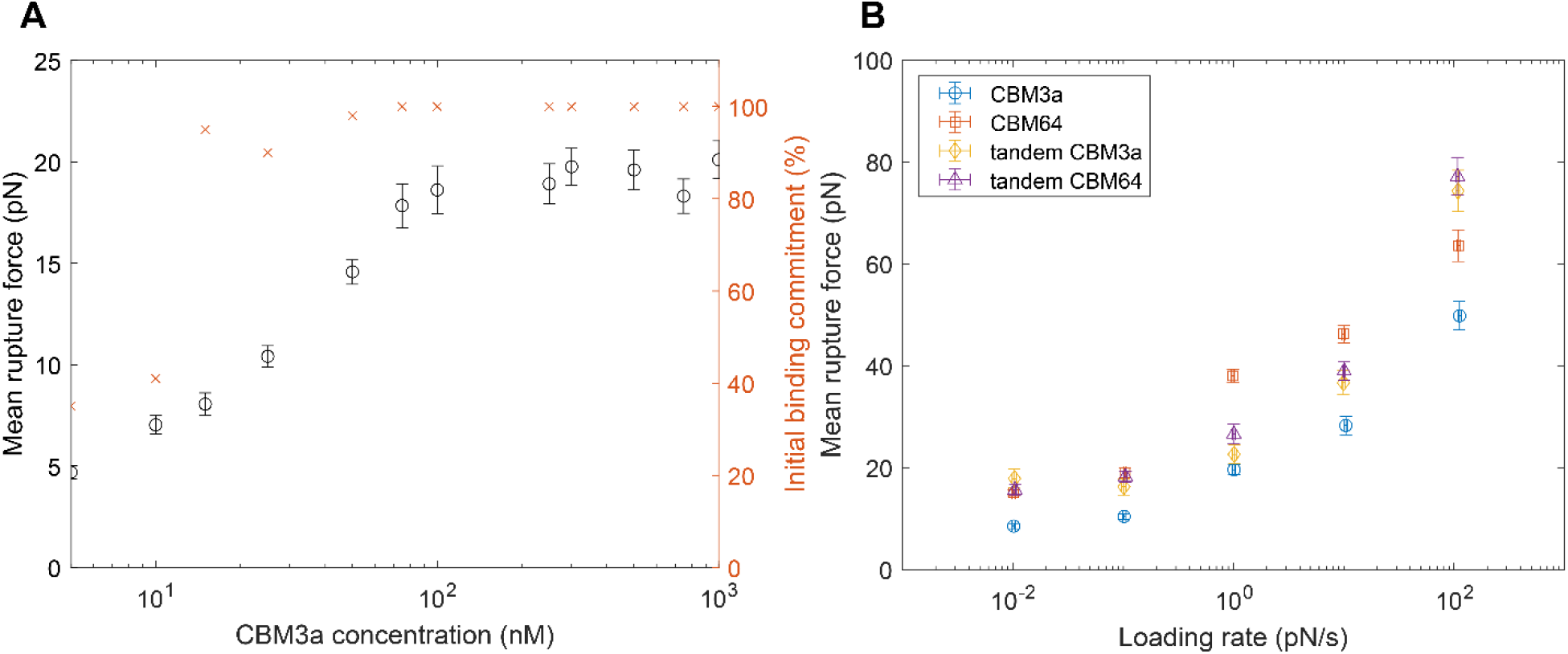
Effect of CBM concentration and loading rate on the mean rupture force of effector beads. A) Beads functionalized with varying concentration of CBM3a at a loading rate of 1 pN/s display no change in the rupture force for concentrations > 100 nM, indicating that the number of CBMs interacting with the surface at once is limited. However, decreasing the concentration below 100 nM reveals a strong effect on the rupture force. Similarly, the initial binding commitment drops significantly below 15 nM, indicating that there is a critical number of CBMs on the bead needed for a successful binding event within the set incubation time of 15 minutes. B) Mean rupture force vs. loading rate for CBM3a, CBM64 and tandem constructs at 500 nM CBM concentration, respectively. While CBM3a-functionalized beads display the lowest rupture force, the difference between CBM constructs is small for loading rates < 0.1 pN/s and spreads as the loading rate increases. Error bars in A) and B) represent the 95% CI of the normal and lognormal histogram fits.

One reason for a higher initial binding commitment despite a drop in rupture force may be due to surface recognition effects associated with the binding rate (k_on_) of CBM3a. Once a single CBM binds to the cellulose surface, other CBMs may bind faster due to proximity effects since multiple CBMs are immobilized to the bead surface. Previous Quartz Crystal Microbalance with Dissipation (QCM-D) analysis of classical bulk protein-ligand binding affinity revealed a relatively high binding rate for CBM3a wild type compared to CBM64 as well as CBM3a and CBM64 mutants (37). The rupture force, on the other hand, is predominately influenced by the unbinding rate, k_off_, for a single, reversible bond as described by Bell (55). Once a force is applied, the distance to the cellulose surface for some individual CBM molecules may be too large to facilitate re-binding. Thus, the bead’s rupture force is mainly a function of the remaining CBMs and their unbinding rate. However, compared to previous single-molecule assays, the mean rupture force is still much larger, most likely due to cooperativity (avidity) effects between individual CBMs during force application.

#### Loading rate dependence and binding percentage of effector-functionalized beads

**Figure 4**-B shows the mean of the lognormal fit of rupture force histograms for each CBM construct and the normal fit of the loading rate. Regardless of loading rate, CBM3a-effector beads display the lowest mean rupture force compared to other CBM constructs. At loading rates below 0.1 pN/s, the difference in rupture forces between tandem CBMs and CBM64 is small but increases at higher loading rates. The mean rupture forces for tandem CBMs are in between the range for single CBMs below 10 pN/s, while the tandem CBM rupture forces are higher at 100 pN/s. The loading rate dependence of the rupture force is non-linear for all CBM constructs, consistent with previous single-molecule experiments of CBM3a and CBM1, where it was shown that the bond lifetime and rupture force do not follow a single exponential decay function (36, 56, 57). The individual rupture force histograms for all loading rates are summarized in Supplemental Figure S5.

**Figure 5**-A demonstrates that there is a noticeable difference between the percentage of bound beads following force application for single and tandem CBM functionalized beads. This difference is not readily apparent when solely analyzing the rupture forces. At loading rates up to 1 pN/s, about half of the tandem CBM effector beads remains bound, while more than 80% of single CBM effector beads detach. The fraction of tandem CBM beads that detached, showed similar rupture forces as shown in **Figure 4**-B, indicating a heterogeneous population of tandem CBM functionalized beads. This heterogeneity may be attributed to the binding kinetics as determined with classical solid state or pull down depletion binding assays (38). While CBM3a was shows to bind reversibly in that study, it was demonstrated that CBM64 binds irreversibly to crystalline cellulose I allomorph, however both tandem constructs showed reversible binding for cellulose I. Tandem CBM64 was shown to bind irreversibly to phosphoric acid-swollen cellulose (PASC) or amorphous cellulose only, thus the beads which remain bound in our assay are likely CBMs binding to disordered or amorphous regions of the applied nanocellulose film.

**Figure 5:**
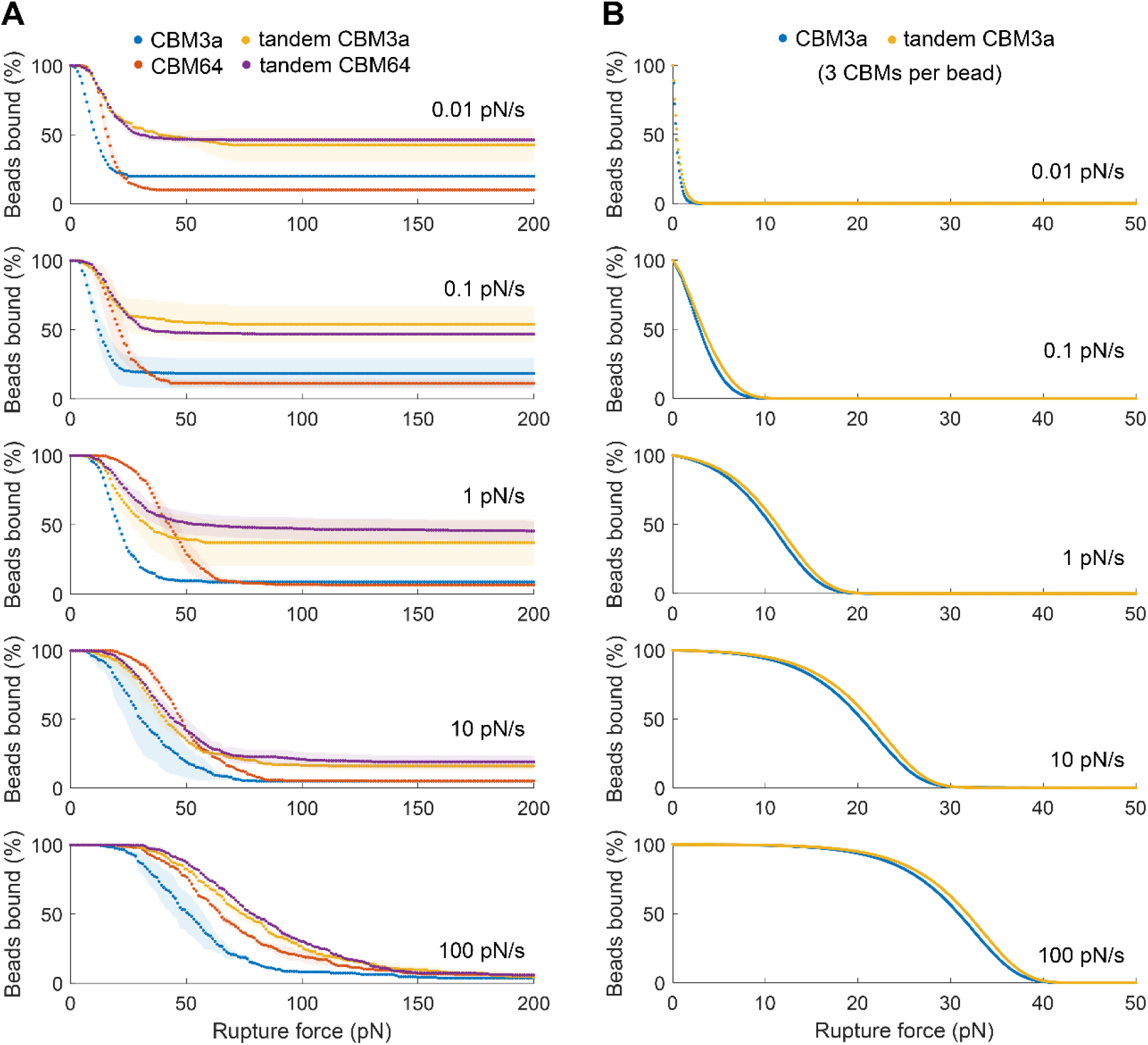
Bound effector beads as a function of the applied force for various loading rates from A) experiments and B) simulations. Regardless of loading rate, the majority of single CBM-functionalized beads detach and apparent avidity effects for tandem CBM-functionalized beads diminish with higher loading rates. A) Around half of tandem CBM functionalized beads remain bound at loading rates up to 1 pN/s and then gradually detach at higher loading rates until at 100 pN/s, all CBM-functionalized beads fully detach at the applied force. The shaded area represents the average deviation of 3 experimental replicates. B) Simulation of the unbinding behavior of a simplified model with 3 CBMs attached to one bead as a function of the loading rate under the no-rebind condition (see Supplemental Information). The 3 CBMs are either a single CBM (blue) or tandem CBM (yellow). Qualitatively, the same trend is observed for both simulations and experiments.

Alternatively, induced avidity effects due to CBMs being bound on beads may enhance their predisposition to irreversibly bind, which cannot be identified using classical pull-down assays. Thus, our assay highlights another novel approach to test reversible protein-ligand binding. As the loading rate increases, the differences in the percentage of bound beads between tandem and single CBM effector beads become less apparent, demonstrating that avidity effects change as a function of the rate at which external forces are applied. Such a behavior may be found relevant to other avidity-based binding systems such as cluster of differentiation (CD) specific CAR T cell binding. The capability of acoustic force-based avidity assays to apply low loading rates which closely match a physiologically relevant shear force regimes is a viable alternative to AFM-based methods (17, 58), where the lowest possible loading rates are in the order of 100 pN/s or higher.

### Modeling the multivalent unbinding of CBM-functionalized beads

To model the observed differences in binding of single and tandem CBM-functionalized beads to cellulose deposited on an AFS chip surface, we first calculated the enhancement of binding for tandem CBM domains connected by a linker relative to that for a single CBM domain. The nucleotide sequence for this linker (see Supplemental Table S1) corresponds to a peptide with 45 residues (GLNATPTKGATPTNTATPTKSATATPTRPSVPTNTPTNTPANTLK) that is predicted to be fully disordered by the disorder prediction servers IUPred3 (59), PONDR-FIT (60), and MobiDB-lite via InterPro (61, 62). We thus modeled the inter-CBM linker as a worm-like chain (WLC) polymer.

**Table 1** shows the predicted effective equilibrium dissociation constants (K_D_’s) for the binding of tandem CBM domains, connected by the disordered linker, to cellulose polymers using the corrected WLC polymer model. In this WLC model, K_D,tandem_ was estimated to be stronger than K_D,single_ for both CBMs. As expected, varying the length of the disordered linker between both CBMs in the tandem construct alters the predicted binding enhancements (Supplemental Table S4) with shorter (or longer) linkers yielding stronger (or weaker) enhancements due to larger (or smaller) C_eff_ factors for the unbound CBM in a single-bound tandem construct.

**Table 1:** Calculated effective dissociation constants for the binding of tandem CBM3a and CBM64 domains to cellulose I using a WLC polymer model of multivalency. The dissociation constants for single CBM constructs were taken from (63). The tandem binding enhancement is calculated as the reciprocal of K_D,tandem_/K_D,single_ as smaller K_D_ values indicate stronger binding.

| System | $K_{D,single}$<br>(μM) | $K_{D,tandem}$<br>(nM) | tandem binding<br>enhancement |
| --- | --- | --- | --- |
| CBM3a | 0.28 | 0.56 | 502x |
| CBM64 | 0.29 | 0.60 | 485x |

We next performed a benchmarking of BioNetGen to determine how many CBM constructs per bead can be handled in our model using a standard workstation. As the dissociation constants for binding to cellulose I are very similar for CBM3a and CBM64 (**Table 1**), we chose to focus on CBM3a in the present simulations to showcase proof-of-concept results. The simulations become very memory intensive with only a small increase in the number of CBMs per bead. The amount of RAM used in the simulations rose exponentially from 1 CBM per bead to 4 CBM per bead (Supplemental Figure S6). Simulations of 4 single CBMs per bead required around 7 GB of RAM to set up all the required reactions from the reaction rules, while only 3 tandem CBMs per bead (equivalent to 6 single CBMs per bead) required around 13GB of RAM. As such, the simulation results described here for both single constructs and tandem constructs are limited to models with 3 CBMs per bead.

Although the number of simulated CBMs per bead is smaller than the hundreds or thousands that can be expected in our experiments, we found that the systems model qualitatively captures the typical behavior of some important experimental trends as seen in **Figure 5**. First, we observed that for the same number of CBMs per bead, larger rupture forces are needed to detach tandem CBM constructs compared to single CBM constructs. As expected, the difference in needed rupture force between tandem and single constructs increased as the number of CBMs per bead increased (highlighted columns in Supplemental Table S5, Supplemental Figure S7) since the tandem constructs have twice as many CBM moieties. All the simulations showed 0% of beads remaining bound to cellulose before the rupture force reached 50 pN (**Figure 5**-B), unlike the experiments which exhibited asymptotic behavior above 0% especially at lower loading rates (**Figure 5**-A). This behavior may likely be due to the much larger number of CBMs per bead attainable in the experiments but not in our simplified simulations. Note that increasing the binding affinity of the beads to cellulose in the simulations either by increasing the kinetic on-rate (k_on_) or decreasing the kinetic off-rate (k_off_) does not capture the observed experimental asymptotic behavior (Supplemental Figures S8 and S9). Second, we observed a rightward shift of the simulation curves to larger rupture forces with increased loading rates as shown in **Figure 5**-B, consistent with the observed experimental trends in in **Figure 5**-A. This is likely due to the constraint that previously bound beads that have detached from cellulose are not allowed to rebind, as simulations where rebinding of detached beads is allowed, show a less pronounced rightward shift to larger rupture forces as the loading rate is increased (Supplemental Figure S10). Interestingly, the difference between the single and tandem curves for a given number of CBMs per bead was found to increase with increased loading rate when detached beads are not allowed to rebind but showed a decrease with increased loading rate when rebinding of detached beads is permissible (compare highlighted columns in Supplemental Tables S5 and S6).

## Conclusion

We demonstrate a method to site-specifically biotinylate proteins in *E. coli in vivo* and purify them using standard metal affinity chromatography. The stable nature of the biotin-streptavidin bond in the presence of imidazole makes it unnecessary to buffer-exchange the proteins for subsequent bead functionalization (64). This represents a significant advantage over *in vitro* biotinylation, which requires one buffer exchange step into suitable buffer for biotinylation and a second purification/ buffer exchange step to remove excess biotin and BirA (65). This convenient *in vivo* biotinylation method could be applied to express a library of protein variants in microplate-based cell cultures (50), with the potential to automate the purification process using suitable liquid handling systems (66). The purified protein variants could then directly be combined with streptavidin-coated beads for screening or characterization using a highly multiplexed AFS or other bead-based assays. In vivo biotinylation of proteins has also been successfully demonstrated in insect and mammalian cells (67, 68), suggesting that our methodology could be expanded to include effector proteins more relevant to cell signaling such as cluster of differentiations or monoclonal antibodies that are challenging to produce in bacterial expression systems.

In the example shown in **Figure 4**-A, a low concentration of effector protein is sufficient to generate a stable rupture force response. As a result, it may not be necessary to account for differences in protein concentration when working with a protein library in microplate-based cell cultures, as long as a minimum amount of effector protein is present. Alternatively, expressing the effector proteins as GFP fusion constructs can provide a simple way to determine the exact protein concentration, if necessary. The process of creating effector beads functionalized with a specific protein is not limited to the biotin-streptavidin interaction. Our workflow can be expanded to also create a covalent bond using alternative systems, such as the SpyCatcher/SpyTag system (69), SNAP tag (70) or by incorporating unnatural amino acids for site specific covalent binding to appropriate receptor beads (e.g., Click Chemistry) (71).

The demonstrated proof-of-concept workflow for preparing effector beads as model systems to investigate cell avidity, can be performed with magnetic tweezers (using magnetic streptavidin-coated beads) or z-Movi® or AFS tweezers (using fluorescently labeled beads), broadening the scope of possible applications. Compared to the z-Movi® technology, it is not necessary to use fluorescently labeled beads (29), since our developed GUI detects a rupture event based on the calibrated z position using well-established methods (72).

Modeling the binding of multiple CBMs (both as tandem and single constructs) linked by a bead yielded valuable insights into the molecular-level interactions between CBM and cellulose ligands. The observed similarities between experimental and simulated trends, as illustrated in **Figure 5**, also support the validity of our model effector bead assay. Moreover, the experimental data can be used to refine model parameters, enhancing our understanding of avidity effects.

Lastly, our findings suggest that both the surface density of effector proteins on beads and the force loading rate can affect the observed rupture force, as well as the percentage of bound beads that remain after force application. These factors may potentially impact the conclusions drawn from force-based avidity assays. Our results emphasize the importance of considering these parameters when designing future experiments to investigate cell avidity using effector beads or cells. Ultimately, such experimental assays may contribute to a better understanding of multivalent cell-cell or cell-surface binding interactions in the presence of external forces.

## Acknowledgements

SPSC acknowledges funding support from the National Science Foundation (CBET Career Award 1846797) and Rutgers Aresty Program. M.H. would like to thank Prof. Timo Betz for kindly sharing the Kitsune source code and Ms. Shriya Kuruba for her help with the initial version of the analysis software. M.H. further thanks Mrs. Karina Hackl for beta-testing the GUI. Figure 1 was created using Biorender.com.

## References

1. G. Carrard, A. Koivula, H. Söderlund, P. Béguin, Cellulose-binding domains promote hydrolysis of different sites on crystalline cellulose. Proc. Natl. Acad. Sci. U. S. A. 97, 10342–10347 (2000).

2. R. Brunecky, et al., Synthetic fungal multifunctional cellulases for enhanced biomass conversion. Green Chem. 22, 478–489 (2020).

3. K. Zajki-Zechmeister, G. S. Kaira, M. Eibinger, K. Seelich, B. Nidetzky, Processive enzymes kept on a leash: How cellulase activity in multienzyme complexes directs nanoscale deconstruction of cellulose. ACS Catal. 11, 13530–13542 (2021).

4. Y. Lu, Y. H. P. Zhang, L. R. Lynd, Enzyme-microbe synergy during cellulose hydrolysis by Clostridium thermocellum. Proc. Natl. Acad. Sci. U. S. A. 103, 16165–16169 (2006).

5. S. P. S. Chundawat, et al., Saccharification of thermochemically pretreated cellulosic biomass using native and engineered cellulosomal enzyme systems. React. Chem. Eng. 1, 616–628 (2016).

6. R. O. Hynes, Integrins: Versatility, modulation, and signaling in cell adhesion. Cell 69, 11–25 (1992).

7. K. Ley, C. Laudanna, M. I. Cybulsky, S. Nourshargh, Getting to the site of inflammation: The leukocyte adhesion cascade updated. Nat. Rev. Immunol. 7, 678–689 (2007).

8. R. Alon, K. Ley, Cells on the run: shear-regulated integrin activation in leukocyte rolling and arrest on endothelial cells. Curr. Opin. Cell Biol. 20, 525–532 (2008).

9. S. Erlendsson, K. Teilum, Binding Revisited—Avidity in Cellular Function and Signaling. Front. Mol. Biosci. 7, 1–13 (2021).

10. V. A. Correa, T. S. Rodrigues, A. I. Portilho, G. Trzewikoswki de Lima, E. De Gaspari, Modified ELISA for antibody avidity evaluation: The need for standardization. Biomed. J. 44, 433–438 (2021).

11. C. H. June, R. S. O’Connor, O. U. Kawalekar, S. Ghassemi, M. C. Milone, CAR T cell immunotherapy for human cancer. Science (80-.). 359, 1361–1365 (2018).

12. T. Bald, M. F. Krummel, M. J. Smyth, K. C. Barry, The NK cell–cancer cycle: advances and new challenges in NK cell–based immunotherapies. Nat. Immunol. 21, 835–847 (2020).

13. S. Viganò, et al., Functional avidity: A measure to predict the efficacy of effector T cells? Clin. Dev. Immunol. 2012 (2012).

14. P. I. Kitov, D. R. Bundle, On the Nature of the Multivalency Effect: A Thermodynamic Model. J. Am. Chem. Soc. 125, 16271–16284 (2003).

15. K. M. Müller, K. M. Arndt, A. Plückthun, Model and simulation of multivalent binding to fixed ligands. Anal. Biochem. 261, 149–158 (1998).

16. W. J. Errington, B. Bruncsics, C. A. Sarkar, Mechanisms of noncanonical binding dynamics in multivalent protein–protein interactions. Proc. Natl. Acad. Sci. U. S. A. 116, 25659–25667 (2019).

17. M. Benoit, D. Gabriel, G. Gerisch, H. E. Gaub, Discrete interactions in cell adhesion measured by single-molecule force spectroscopy. Nat. Cell Biol. 2, 313–317 (2000).

18. C. B. Gilfillan, M. Hebeisen, N. Rufer, D. E. Speiser, Constant regulation for stable CD8 T-cell functional avidity and its possible implications for cancer immunotherapy. Eur. J. Immunol. 51, 1348–1360 (2021).

19. R. B. M. Schasfoort, F. Abali, I. Stojanovic, G. Vidarsson, L. W. M. M. Terstappen, Trends in SPR cytometry: Advances in label-free detection of cell parameters. Biosensors 8, 1–11 (2018).

20. H. Lu, et al., Microfluidic shear devices for quantitative analysis of cell adhesion. Anal. Chem. 76, 5257–5264 (2004).

21. 1. A. J. García, N. D. Gallant, Stick and grip: Measurement systems and quantitative analyses of integrin-mediated cell adhesion strength. Cell Biochem. Biophys. 39, 61–73 (2003).

22. J. C. Friedland, M. H. Lee, D. Boettiger, Mechanically Activated Integrin Switch Controls alpha5 beta1 function. Science (80-.). 323, 642–644 (2009).

23. D. Kamsma, et al., Single-Cell Acoustic Force Spectroscopy: Resolving Kinetics and Strength of T Cell Adhesion to Fibronectin. Cell Rep. 24, 3008–3016 (2018).

24. A. Katsarou, et al., Combining a CAR and a chimeric costimulatory receptor enhances T cell sensitivity to low antigen density and promotes persistence. Sci. Transl. Med. 13, 1–17 (2021).

25. R. C. Larson, et al., CAR T cell killing requires the IFNγR pathway in solid but not liquid tumours. Nature 604, 563–570 (2022).

26. N. Balneger, et al., Sialic acid blockade in dendritic cells enhances CD8+ T cell responses by facilitating high-avidity interactions. Cell. Mol. Life Sci. 79, 1–15 (2022).

27. L. Halim, et al., Engineering of an Avidity-Optimized CD19-Specific Parallel Chimeric Antigen Receptor That Delivers Dual CD28 and 4-1BB Co-Stimulation. Front. Immunol. 13, 1–13 (2022).

28. M. B. Leick, et al., Non-cleavable hinge enhances avidity and expansion of CAR-T cells for acute myeloid leukemia. Cancer Cell 40, 494-508.e5 (2022).

29. Y. Wang, J. Jin, H. J. Wang, L. A. Ju, Acoustic Force-Based Cell–Matrix Avidity Measurement in High Throughput. Biosensors 13, 95 (2023).

30. T. Buranda, et al., Rapid parallel flow cytometry assays of active GTPases using effector beads. Anal. Biochem. 442, 149–157 (2013).

31. S. L. Schwartz, et al., Flow cytometry for real-time measurement of guanine nucleotide binding and exchange by Ras-like GTPases. Anal. Biochem. 381, 258–266 (2008).

32. Y. Li, R. J. Kurlander, Comparison of anti-CD3 and anti-CD28-coated beads with soluble anti-CD3 for expanding human T cells: Differing impact on CD8 T cell phenotype and responsiveness to restimulation. J. Transl. Med. 8, 104 (2010).

33. W. C. Chan, Y. C. Linn, A comparison between cytokine- and bead-stimulated polyclonal T cells: the superiority of each and their possible complementary role. Cytotechnology 68, 735–748 (2016).

34. P. J. Schatz, Use of Peptide Libraries to Map the Substrate Specificity of a Peptide-Modifying Enzyme: A 13 Residue Consensus Peptide Specifies Biotinylation in Escherichia coli. Nat. Biotechnol. 11, 1138–1143 (1993).

35. R. S. Reiner, A. W. Rudie, “Process scale-up of cellulose nanocrystal production to 25 kg per batch at the Forest Products Laboratory” in Production and Applications of Cellulose Nanomaterials, (TAPPI Press, 2013), pp. 21–24.

36. M. Hackl, et al., Acoustic force spectroscopy reveals subtle differences in cellulose unbinding behavior of carbohydrate-binding modules. Proc. Natl. Acad. Sci. 119, e2117467119 (2022).

37. B. Nemmaru, et al., Reduced type-A carbohydrate-binding module interactions to cellulose I leads to improved endocellulase activity. Biotechnol. Bioeng. 118, 1141–1151 (2021).

38. D. Jayachandran, et al., Engineering and characterization of carbohydrate-binding modules to enable real-time imaging of cellulose fibrils biosynthesis in plant protoplasts. bioRxiv (2023) 10.1101/2023.01.02.522519.

39. M. Z. Li, S. J. Elledge, “SLIC: A Method for Sequence- and Ligation-Independent Cloning” in (2012), pp. 51–59.

40. M. Fairhead, M. Howarth, “Site-Specific Biotinylation of Purified Proteins Using BirA” in Site-Specific Protein Labeling: Methods and Protocols, (2015), pp. 171–184.

41. G. Sitters, et al., Acoustic force spectroscopy. Nat. Methods 12, 47–50 (2014).

42. A. Nguyen, M. Brandt, T. M. Muenker, T. Betz, Multi-oscillation microrheology via acoustic force spectroscopy enables frequency-dependent measurements on endothelial cells at high-throughput. Lab Chip, 1929–1947 (2021).

43. D. Kamsma, R. Creyghton, G. Sitters, G. J. L. Wuite, E. J. G. Peterman, Tuning the Music: Acoustic Force Spectroscopy (AFS) 2.0. Methods 105, 26–33 (2016).

44. A. Sethi, B. Goldstein, S. Gnanakaran, Quantifying intramolecular binding in multivalent interactions: A Structure-Based synergistic study on Grb2-Sos1 complex. PLoS Comput. Biol. 7, 1– 13 (2011).

45. T. Travers, et al., Combinatorial diversity of Syk recruitment driven by its multivalent engagement with FcεRIγ. Mol. Biol. Cell 30, 2331–2347 (2019).

46. V. S. Chauhan, S. K. Chakrabarti, Use of nanotechnology for high performance cellulosic and papermaking products. Cellul. Chem. Technol. 46, 389–400 (2012).

47. J. R. Faeder, M. L. Blinov, W. S. Hlavacek, “Rule-Based Modeling of Biochemical Systems with BioNetGen” in Systems Biology, I. V Maly, Ed. (Humana Press, 2009), pp. 113–167.

48. L. A. Harris, et al., BioNetGen 2.2: Advances in rule-based modeling. Bioinformatics 32, 3366– 3368 (2016).

49. Y. Li, R. Sousa, Novel system for in vivo biotinylation and its application to crab antimicrobial protein scygonadin. Biotechnol. Lett. 34, 1629–1635 (2012).

50. B. W. Li, Y. Zhang, Y. C. Wang, Y. Xue, X. Y. Nie, Rapid Fabrication of Protein Microarrays via Autogeneration and on-Chip Purification of Biotinylated Probes. ACS Synth. Biol. 9, 2267–2273 (2020).

51. C. K. Bandi, A. Goncalves, S. V. Pingali, S. P. S. Chundawat, Carbohydrate-binding domains facilitate efficient oligosaccharides synthesis by enhancing mutant catalytic domain transglycosylation activity. Biotechnol. Bioeng. 117, 2944–2956 (2020).

52. P. A. Smith, et al., A plasmid expression system for quantitative in vivo biotinylation of thioredoxin fusion proteins in Escherichia coli. Nucleic Acids Res. 26, 1414–1420 (1998).

53. K. L. Tsao, B. DeBarbieri, H. Michel, D. S. Waugh, A versatile plasmid expression vector for the production of biotinylated proteins by site-specific, enzymatic modification in Escherichia coli. Gene 169, 59–64 (1996).

54. M. Hackl, K. A. Ramdin, S. Chundawat, AFS-GUI (2024) 10.5281/zenodo.1234.

55. G. I. Bell, Models for the specific adhesion of cells to cells. Science (80-.). 200, 618–627 (1978).

56. M. Zhang, B. Wang, B. Xu, Measurements of single molecular affinity interactions between carbohydrate-binding modules and crystalline cellulose fibrils. Phys. Chem. Chem. Phys. 15, 6508–6515 (2013).

57. S. P. S. Chundawat, et al., Molecular origins of reduced activity and binding commitment of processive cellulases and associated carbohydrate-binding proteins to cellulose III. J. Biol. Chem. 296, 100431 (2021).

58. F. Li, S. D. Redick, H. P. Erickson, V. T. Moy, Force measurements of the α5β1 integrin-fibronectin interaction. Biophys. J. 84, 1252–1262 (2003).

59. G. Erdos, M. Pajkos, Z. Dosztányi, IUPred3: Prediction of protein disorder enhanced with unambiguous experimental annotation and visualization of evolutionary conservation. Nucleic Acids Res. 49, W297–W303 (2021).

60. B. Xue, R. L. Dunbrack, R. W. Williams, A. K. Dunker, V. N. Uversky, PONDR-FIT: A meta-predictor of intrinsically disordered amino acids. Biochim. Biophys. Acta - Proteins Proteomics 1804, 996–56. 1010 (2010).

61. M. Necci, D. Piovesan, D. Clementel, Z. Dosztányi, S. C. E. Tosatto, MobiDB-lite 3.0: Fast consensus annotation of intrinsic disorder flavors in proteins. Bioinformatics 36, 5533–5534 (2020).

62. M. Blum, et al., The InterPro protein families and domains database: 20 years on. Nucleic Acids Res. 49, D344–D354 (2021).

63. D. Jayachandran, et al., Engineering and characterization of carbohydrate-binding modules for imaging cellulose fibrils biosynthesis in plant protoplasts. Biotechnol. Bioeng. 120, 2253–2268 (2023).

64. A. Reichel, et al., Noncovalent, site-specific biotinylation of histidine-tagged proteins. Anal. Chem. 79, 8590–8600 (2007).

65. Y. Li, R. Sousa, Expression and purification of E. coli BirA biotin ligase for in vitro biotinylation. Protein Expr. Purif. 82, 162–167 (2012).

66. A. Matte, High-throughput, parallelized and automated protein purification for therapeutic antibody development. Approaches to Purification, Anal. Charact. Antibody-Based Ther., 181–198 (2020).

67. G. Strübbe, et al., Polycomb purification by in vivo biotinylation tagging reveals cohesin and Trithorax group proteins as interaction partners. Proc. Natl. Acad. Sci. U. S. A. 108, 5572–5577 (2011).

68. E. de Boer, et al., Efficient biotinylation and single-step purification of tagged transcription factors in mammalian cells and transgenic mice. Proc. Natl. Acad. Sci. 100, 7480–7485 (2003).

69. B. Zakeri, M. Howarth, Spontaneous intermolecular amide bond formation between side chains for irreversible peptide targeting. J. Am. Chem. Soc. 132, 4526–4527 (2010).

70. R. Dreyer, R. Pfukwa, S. Barth, R. Hunter, B. Klumperman, The Evolution of SNAP-Tag Labels. Biomacromolecules (2022) 10.1021/acs.biomac.2c01238.

71. A. Adhikari, et al., Reprogramming natural proteins using unnatural amino acids. RSC Adv. 11, 38126–38145 (2021).

72. M. T. J. Van Loenhout, J. W. J. Kerssemakers, I. De Vlaminck, C. Dekker, Non-bias-limited tracking of spherical particles, enabling nanometer resolution at low magnification. Biophys. J. 102, 2362–2371 (2012).

