## Supplementary Information for "Site-specific Effector Protein Functionalization to Create Bead-based Avidity Model Systems"

**This SI includes:**

Supplemental Methods

Supplemental Figures S1 to S10

Supplemental Table S1 to S6

Supplemental Information References

Supplemental Files 1 and 2 (BioNetGen input files for single and tandem CBM modeling)

#### Supplemental methods

**Modeling of multivalent protein-ligand binding**

***Worm-like chain (WLC) model of the disordered linker in tandem CBM constructs***

For a molecule containing two binding domains with equilibrium association constants of $K_{A,1}$ and $K_{A,2}$, and that are separated by a linker, the effective equilibrium association constant ($K_{A,overall}$) for binding of both domains to a molecule with multiple ligand sites is given by equation S1

$K_{A,overall}=C_{eff}\times K_{A,1}\times K_{A,2}$	(S1)

where, after ligand binding of one of the two domains, $C_{eff}$ represents the effective concentration of unbound ligand experienced by the unbound domain (after binding of the other domain). It was shown previously (1, 2) that if the distance distribution between the two binding domains is given by $p_{D-D}\left( r \right)$ and the corresponding distribution for ligand sites is $p_{L-L}\left( r \right)$, then $C_{eff}$ can be calculated by finding the overlapping regions between the two distributions and integrating the products of the probabilities in these regions for all possible distances as given in equation S2:

$C_{eff}=\int_{r=0}^{\infty} p_{D-D}\left( r \right) p_{L-L}\left( r \right) d^{3}r$	(S2)

For the binding of tandem CBM domains to cellulose polymers, we employed a worm-like chain (WLC) polymer model to get the distance distribution $p_{D-D}\left( r \right)$ by assuming that the linker between both CBM domains is fully disordered. The WLC model derivation for this distribution yields equation S3:

$p_{D-D}\left( r \right)=\left( \frac{3}{4\pi l_{p}l_{c}} \right)^{\frac{3}{2}}\times\exp\left( \frac{-3r^{2}}{4l_{p}l_{c}} \right)$	(S3)

where $l_{p}$ is the linker’s persistence length that gives the end-to-end length beyond which the linker behaves as a random walk rather than a flexible rod (3) as described in equation S4:

$l_{p}=0.074\times N^{0.43}$	(S4)

and $l_{c}$ is the linker’s contour length that gives its maximum end-to-end length when fully extended (4) in equation S5:

$l_{c}=0.38\times N$	(S5)

In equations S4 and S5, $N$ is the number of residues in the disordered linker. For the linker between the two CBM domains, $N=45$. Shorter ($N=10$) and longer ($N=100$) linkers were also tested to assess the impact of linker size on tandem binding affinity.

As the cellulose polymers are attached to the surface of the AFS chip, we assumed a uniform distance distribution $p_{L-L}\left( r \right)$ for distances up to the CBM linker’s contour length $l_{c}$ (*i.e.*, the maximum possible distance between tandem CBM domains). $C_{eff}$ is then calculated with equation S2 using the WLC-derived distribution $p_{D-D}\left( r \right)$ and the uniform distribution $p_{L-L}\left( r \right)$. Compared with previous work where the ligands are free-floating, the cellulose polymers are restricted here to a surface thereby reducing the rate of productive encounter complex formation for the tandem domain. We thus incorporated here a volume-based corrective factor that reduces the $C_{eff}$ of unbound ligand experienced by the tandem domain as described in equation S6:

$C_{eff,corrected}= C_{eff}\times\frac{V_{slab}}{V_{hemi}}$	(S6)

where $V_{slab}$ is the volume of a slab above the chip that is occupied by attached cellulose polymers, equation S7:

$V_{slab}=4\pi r^{2}h$	(S7)

with $r=2.5 \mu m$ representing the radius of the polystyrene bead to which CBM constructs are functionalized, and $h=6 nm$ giving the average diameter of a cellulose microfibril (5) (used here as the average height of the slab above the chip); while $V_{hemi}$ is the volume of a hemisphere above the chip as defined in equation S8:

$V_{hemi}=\frac{2}{3}\pi r^{3}$	(S8)

with also $r=2.5 \mu m$. Equation S8 gives the search volume of the tandem domain if the ligands are free-floating, while equation S7 gives the search volume when the ligands are restricted to the chip surface. $C_{eff,corrected}$ can then be used in equation S1 to calculate $K_{A,overall}$, whose reciprocal is the effective equilibrium dissociation constant for tandem binding ($K_{D,overall}$). Note that $K_{A,1}=K_{A,2}$ in equation S1 since the tandem constructs here always contain the same CBM domain types (either both CBM3a or both CBM64).

***Network models of cellulose binding by beads functionalized with single or tandem CBM constructs***

Rule-based models were developed to simulate the binding of beads containing either single or tandem CBM constructs to cellulose polymers using BioNetGen v2.9.0 (6, 7). This network modeling software can generate a complete reaction network along with the corresponding ordinary differential equations (ODEs) from a limited number of reaction rules describing the system. The ODEs are then numerically integrated using a CVODE interface provided by BioNetGen to simulate the time-dependent concentrations of all species. Although BioNetGen supports the use of multiple compartments and volumetric scaling of reactions, we used a simplified model here that places all molecules in a single well-mixed compartment. In order to reduce the complexity of the generated reaction networks, we added an additional assumption that all the CBM sites on a single bead can bind only to ligand sites from a single cellulose polymer.

The CBM-functionalized beads are represented in the models by the species described in equation S9:

$B\left( i!1,i!2 \right).Re\left( i!1,l,l \right).Re(i!2,l,l)$	(S9)

where $B$ represents the bead and $Re$ represents the CBM construct or receptor. The CBM constructs are bound to the beads via the $i$ sites, and the number of constructs per bead can be increased by joining more $Re\left( i!\#,l,l \right)$ moieties to the bead (along with a corresponding increase in the number of bound $i$ sites on the bead). Equation S9 describes receptors with tandem CBM domains (*i.e.*, two $l$ sites); receptors with single CBM domains each have a single $l$ site.

Cellulose polymers are represented in both single CBM and tandem CBM models as defined in equation S10:

$L(r,r,r,r,r,r,r,r)$	(S10)

with each $r$ site representing a binding site for a CBM. Note that a single type A CBM domain (such as CBM3a and CBM64) binds to multiple sugar moieties on cellulose; each group of multiple sugar moieties is simplified here as a single site $r$.

Initial binding of a CBM-functionalized bead to cellulose is given by the following reversible reaction rule in equation S11:

$B\left( i!1,i!2 \right).Re\left( i!1,l,l \right).Re\left( i!2,l,l \right)+ L\left( r,r,r,r,r,r,r,r \right)\leftrightarrow B\left( i!1,i!2 \right).Re\left( i!1,l!3,l \right).Re\left( i!2,l,l \right).L\left( r!3,r,r,r,r,r,r,r \right) k_{on},k_{off}$	(S11)

where $k_{on}$ and $k_{off}$ are the association and dissociation rate constants, respectively. Experimentally measured values were used for these rate constants (8, 9). Tandem binding is given by the following reversible reaction rule in equation S12:

$Re\left( l!1,l \right)+ L\left( r!1,r \right)\leftrightarrow Re\left( l!1,l!2 \right).L\left( r!1,r!2 \right) k_{on,tandem},k_{off}$	(S12)

where $k_{off}$ is the same dissociation rate constant, while $k_{on,tandem}$ is the association rate constant tandem and is the product of $k_{on}$ and one of the tandem binding enhancement factors in Table S4. Equation S11 is a longer rule as it describes the initial binding events and so the unbound states of the $l$ and $r$ states need to be explicitly specified. Whereas equation S12 provides a shorter rule since tandem binding of a receptor to cellulose does not depend on the bound/unbound state of the other receptors on the bead.

Each simulation comprised two phases. The first phase involved system equilibration until the concentrations of all species reached steady-state levels. Beads that fully detach from cellulose during this phase are allowed to rebind. In the second phase, a force is added to the system to simulate the application of acoustic force in the experiments. Bell’s model was used to estimate the impact of applied physical force on the dissociation time (and thus the dissociation rate) of biomolecular bonds (10) as described in equation S13:

$k_{off}\left( f \right)= k_{off}^{0}\times\exp\left( \frac{x^{\ddagger}f}{k_{B}T} \right)$	(S13)

where $f$ is the force applied to the system, $k_{off}^{0}$ is the dissociation rate constant in the absence of applied force, $k_{off}\left( f \right)$ is the dissociation rate constant as a function of the applied force, $k_{B}$ is the Boltzmann constant, $T$ is the absolute temperature, and $x^{\ddagger}$ is a parameter that describes the distance between bound and transition states. The latter parameter was obtained from fits of experimental measurements (8). The “setParameter” function of BioNetGen was used to update the value of $k_{off}$ (based on equation S13) at multiple time points in the simulation with increasing amount of force applied. To simulate the effect of different loading rates, the time interval between consecutive time points where $k_{off}$ gets updated is varied (*e.g.*, larger loading rates correspond to shorter time intervals between $k_{off}$ updates). For the second phase of the simulations where loading rates were applied, either a “can-rebind” or “no-rebind” approach was used depending on whether beads that have fully detached from cellulose are allowed to rebind or not, respectively. The “no-rebind” approach captures the dynamics of beads that have fully detached from cellulose in the experiments.

Sample BioNetGen input files for the single CBM and tandem CBM network models are provided in Supplemental Files 1 and 2, respectively.

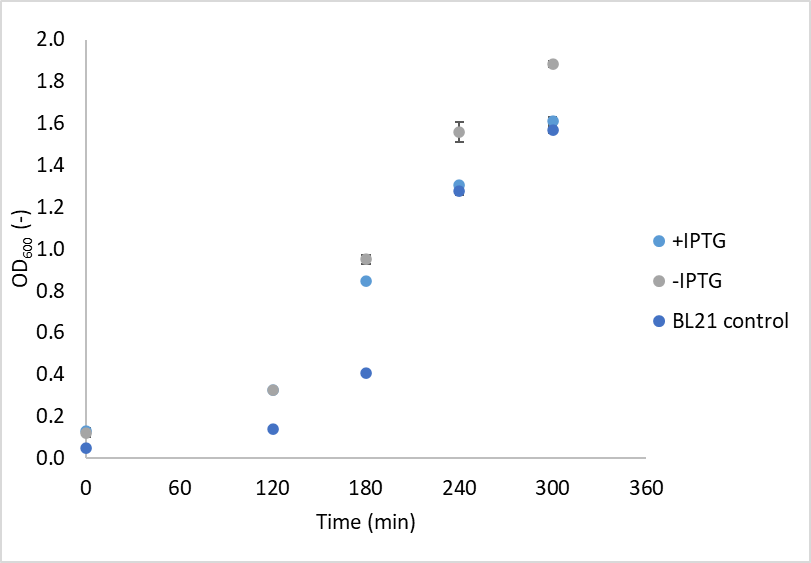

**Supplemental Figure S1:** Growth curve of E. coli BL21 cells harboring the MBP-BirA plasmid in the presence or absence of 1mM IPTG, respectively. The control BL21 does not contain the MBP-BirA plasmid. Cells were grown in 5 ml LB media at 37°C in 50 ml cylindrical tubes with shaking at 200 rpm. The optical density was measured in a clear-bottom 96-well plate using 200 μl of culture and the obtained readings normalized to a path length of 1cm. The error bar represents the average standard deviation from mean of 3 experimental replicates.

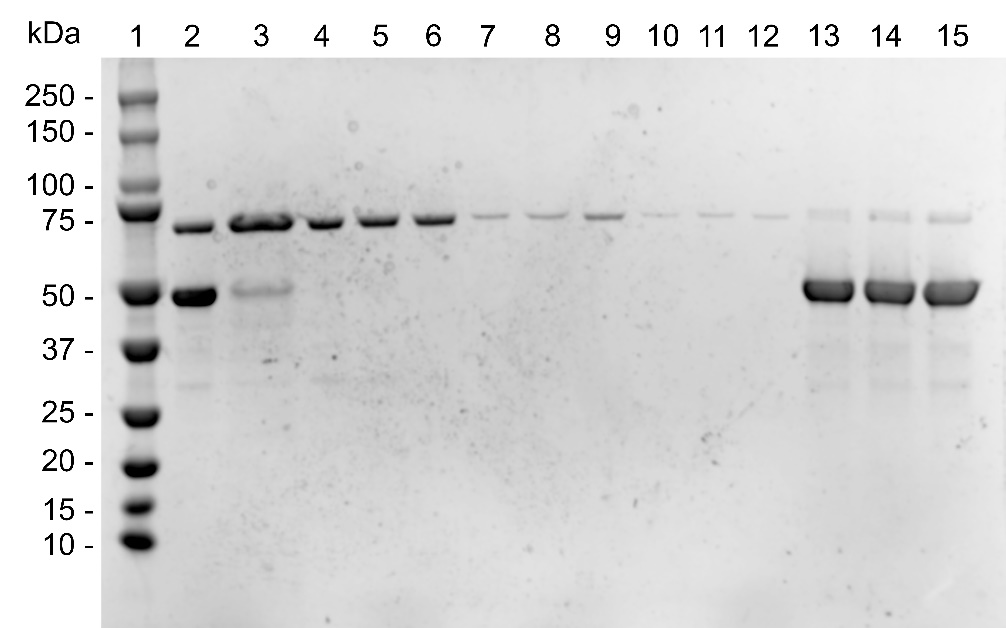

**Supplemental Figure S2:** SDS-PAGE showing the progress of MBP-BirA removal with various buffers and washing steps from IMAC purified GFP-CBM3a fusion protein. Due to the high salt concentration (500mM) and absence of surfactants in the initial lysis and IMAC buffers (refer to main text), BirA formed a stable complex with the Avi-tagged protein, which was removed by subsequent washing steps. The IMAC-purified sample (1 ml) was buffer exchanged into PBS pH 7.4, mixed with 1 ml of Ni-NTA magnetic beads and incubated for 1 hour. After taking a sample for SDS-PAGE, the slurry was split up equally into 3 new 1.5 ml tubes. Next, the beads were incubated 20 minutes each with either PBS (10 mM, pH 7.4) or PBS containing 0.1 % (v/v) Triton-X100 or PEG-4kDa. The incubation and washing steps were repeated twice (3 washing steps in total) and beads were finally eluted in 500 ml of 500 mM imidazole in PBS. Lane 1: protein standard, Lane 2: IMAC purified and buffer exchanged GFP-CBM3a (~56 kDa) contains significant amounts of MBP-BirA (~70kDa), Lane 3: Supernatant after 1 hour of incubation in PBS showing that most histidine-tagged CBM is bound to the beads, Lane 4: Supernatant of PBS+Triton sample after first incubation/wash, Lane 5: Supernatant of PBS+PEG sample after first incubation/wash, Lane 6: Supernatant of PBS sample after first incubation/wash, Lane 7: Supernatant of PBS+Triton sample after second incubation/wash, Lane 8: Supernatant of PBS+PEG sample after second incubation/wash, Lane 9: Supernatant of PBS sample after first incubation/wash, Lane 10: Supernatant of PBS+Triton sample after third incubation/wash, Lane 11: Supernatant of PBS+PEG sample after third incubation/wash, Lane 12: Supernatant of PBS sample after first incubation/wash, Lane 13: Supernatant of PBS+Triton sample after elution in 500mM imidazole, Lane 14: Supernatant of PBS+PEG sample after elution in 500mM imidazole, Lane 15: Supernatant of PBS sample after elution in 500mM imidazole.

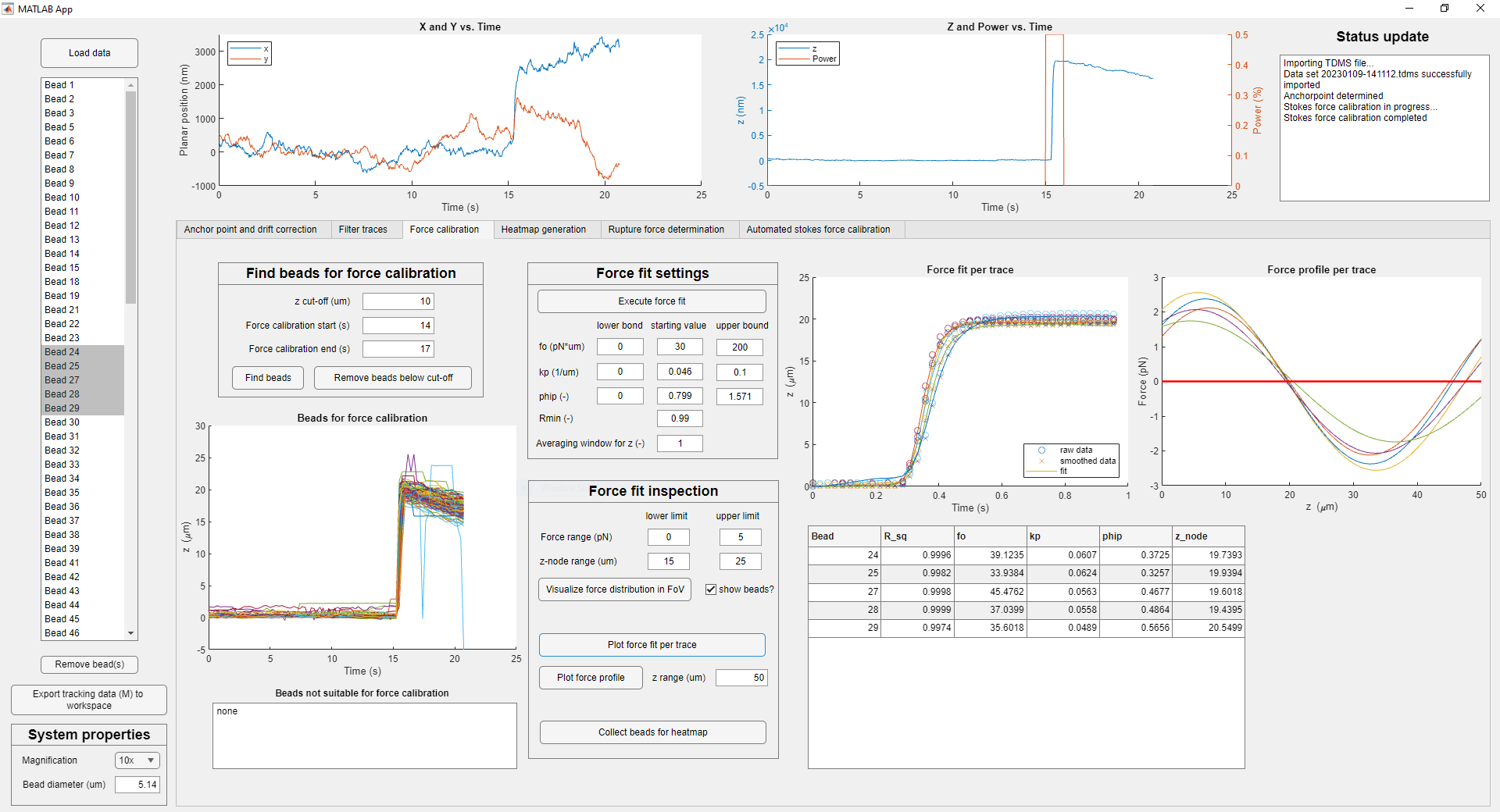

**Supplemental Figure S3:** Screenshot of the developed GUI showing the force calibration tab. After SFC, the force fit as well as force profile of individual traces can be inspected manually. For a detailed description, refer to the GUI manual.

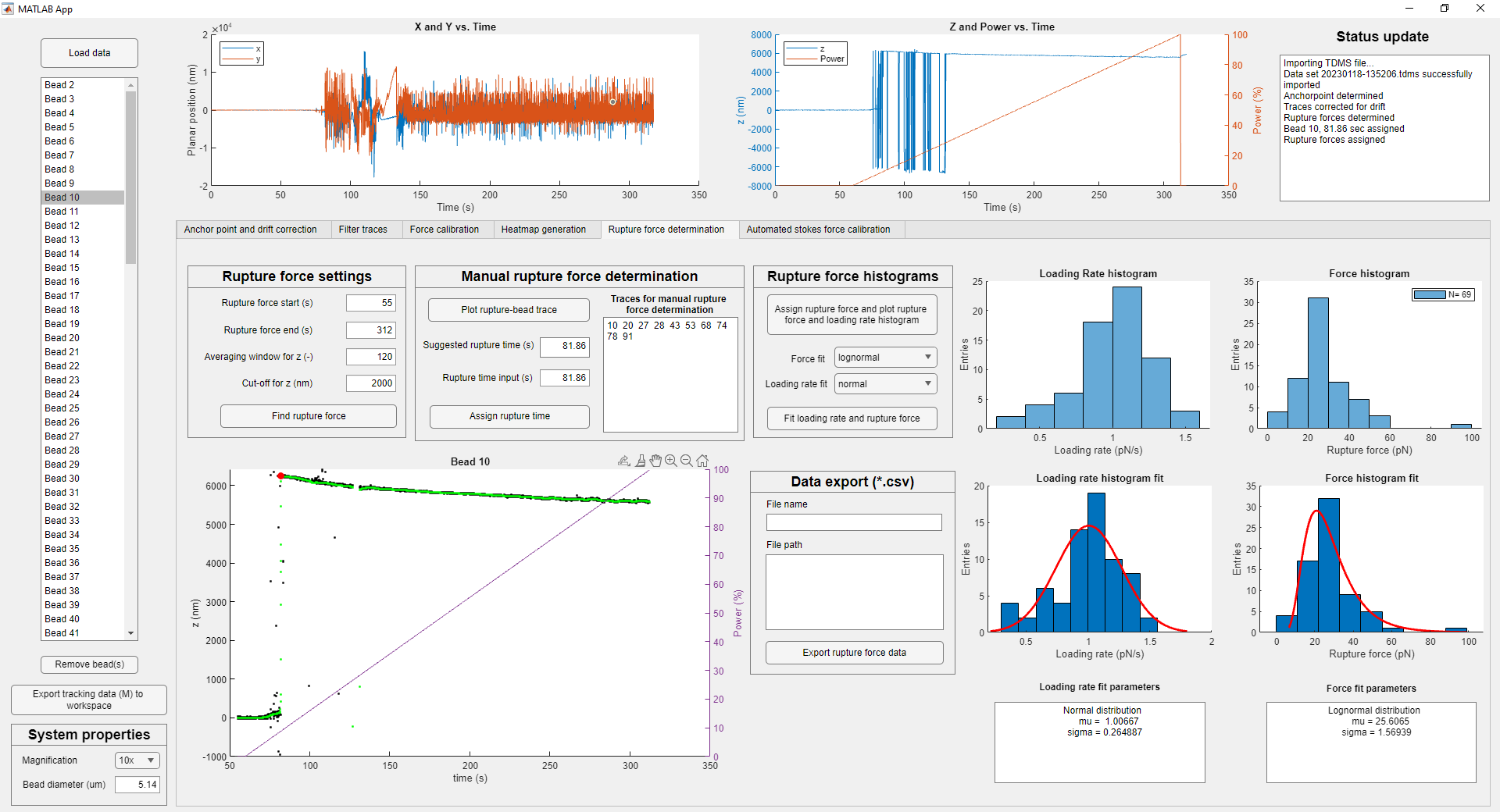

**Supplemental Figure S4:** Screenshot of the developed GUI showing the rupture force determination tab. With a few clicks, the rupture time is accurately determined, and the rupture forces are assigned based on a previously generated force factor heatmap. In addition, the summary of rupture force and loading rate is presented as histograms.

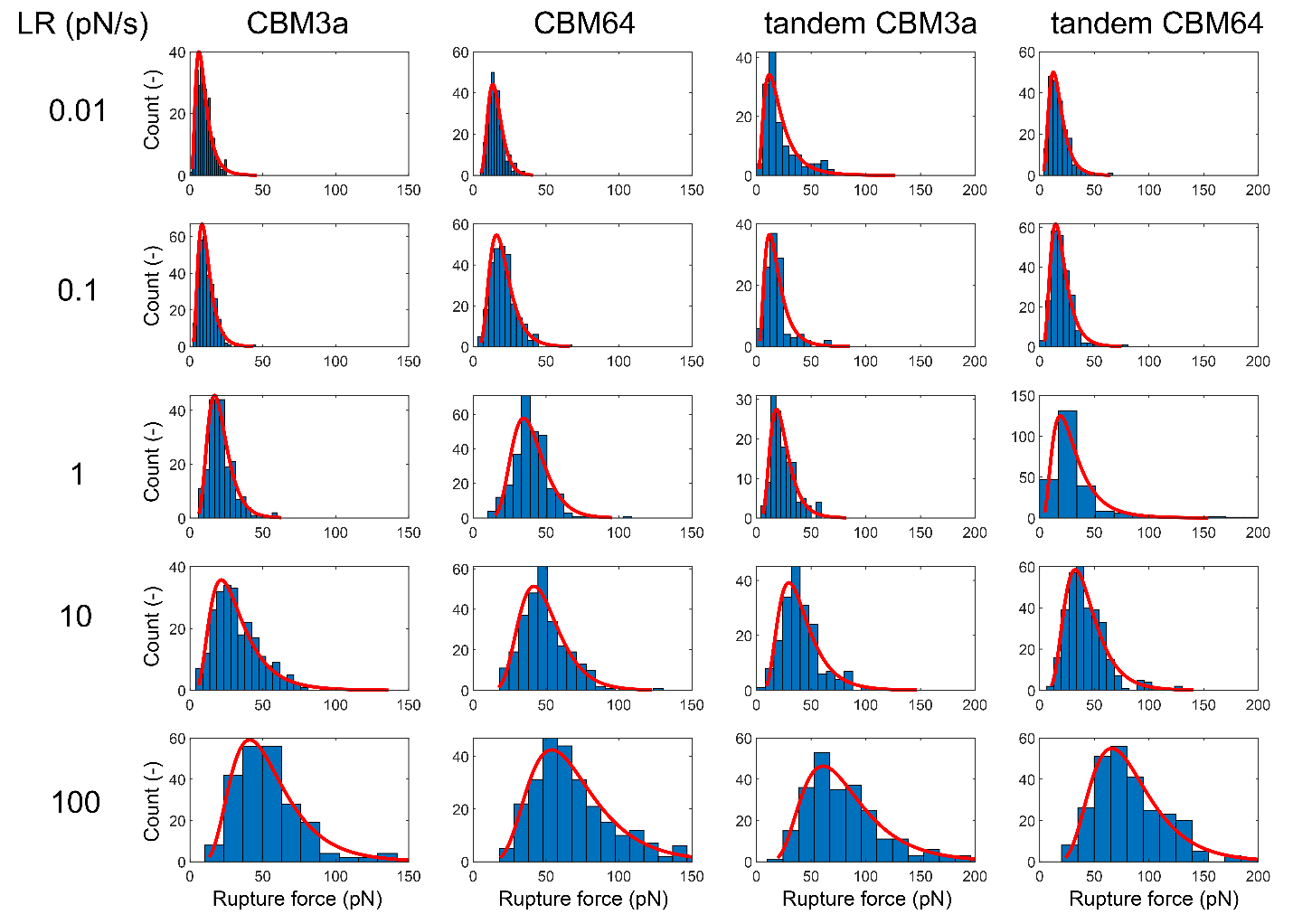

**Supplemental Figure S5**: Rupture force histograms of the 4 tested effector proteins at loading rates (LR) between 0.01 – 100 pN/s.

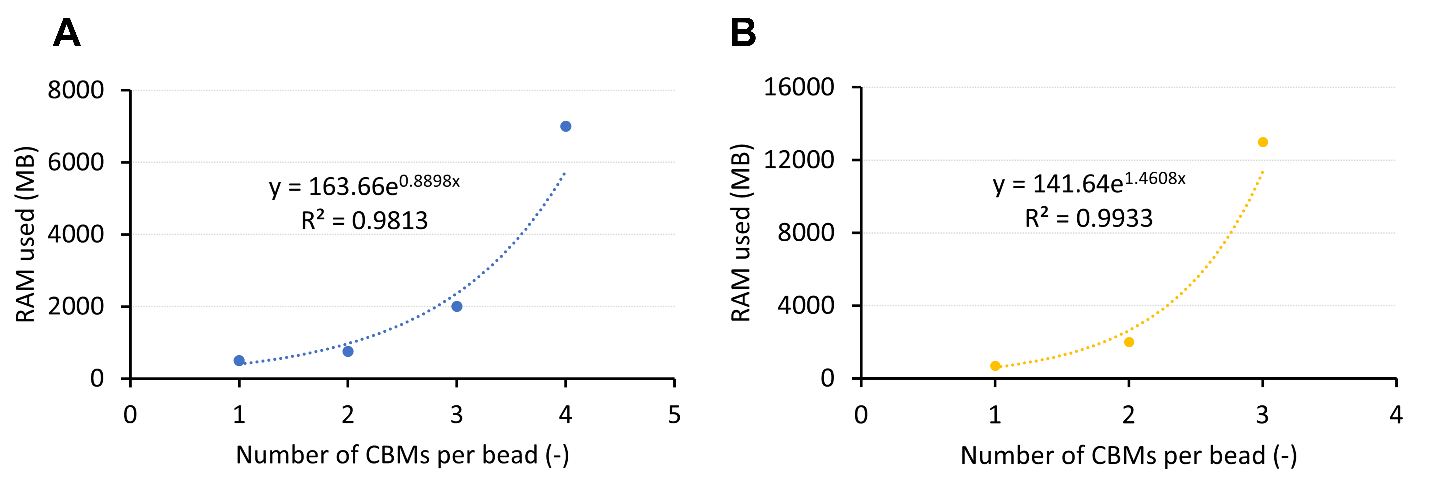

**Supplemental Figure S6**: Benchmarks of amount of RAM used (MB) in BioNetGen simulations of CBM-functionalized beads binding to cellulose polymers as a function of the number of CBM constructs attached to each bead. Data for benchmarks using A) single and B) tandem are shown. The two data sets were fitted with exponential trendlines; the trendline equations and coefficients of determination (R^2^) are provided in both plots. From this fitting, around 14 GB of RAM is needed to simulate the cellulose binding of beads with 5 single CBM constructs each, while around 49 GB of RAM is needed for beads with 4 tandem CBM constructs each.

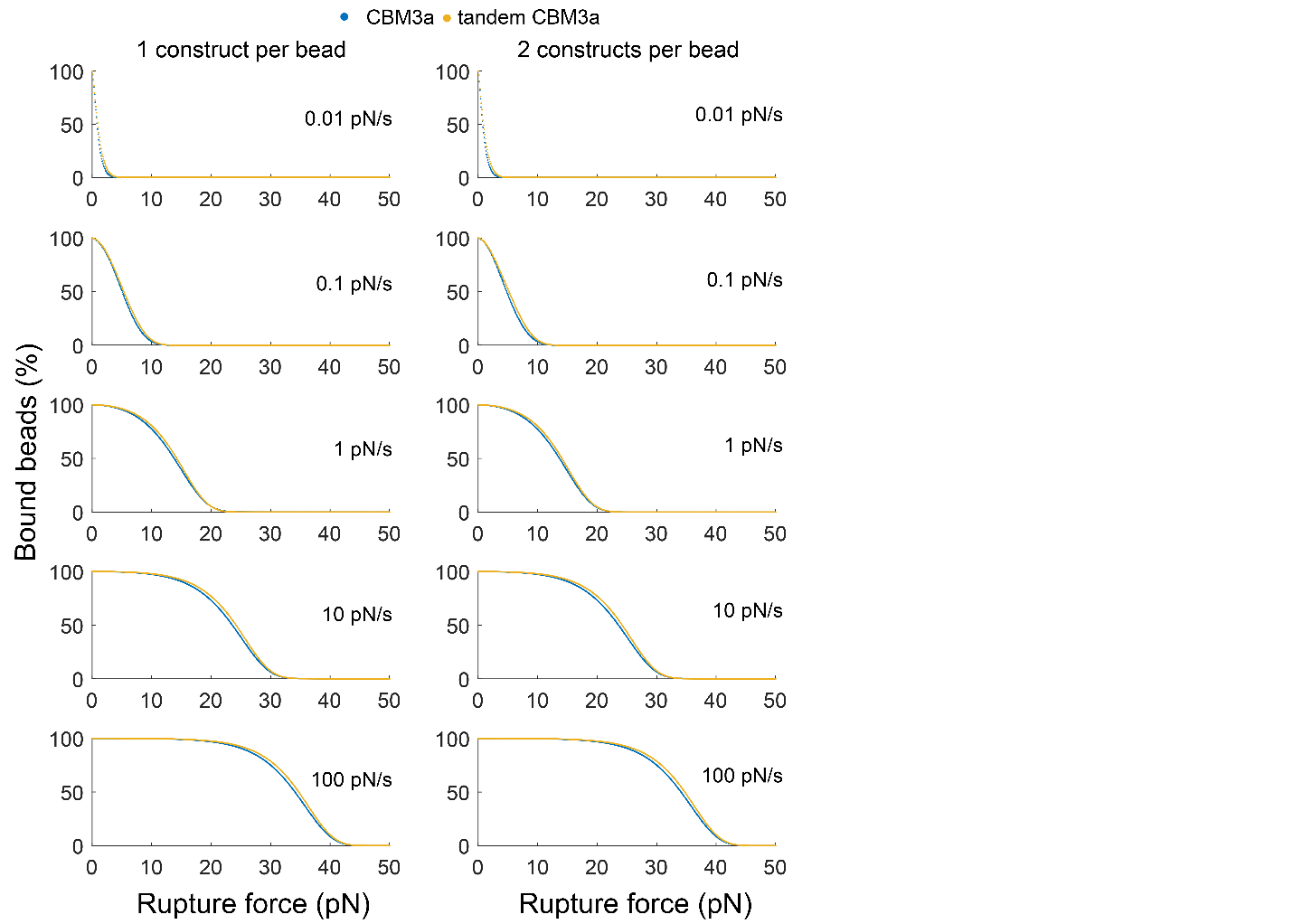

***Supplemental Figure S7****: Simulated percentage of CBM3a-functionalized beads bound to cellulose I polymers as a function of the applied rupture force (pN). Beads are functionalized in the simulations with either single (blue curves) or tandem (yellow curves) CBM constructs. A “no-rebind” WLC polymer model (see Supplemental Methods) was used in these runs where beads that become fully detached from cellulose are not allowed to rebind. Each row gives plots generated for a particular loading rate (from top to bottom: 0.01 pN/s, 0.1 pN/s, 1 pN/s, 10 pN/s, and 100 pN/s). The rupture force in the simulations went up to 200 pN; for visualization, the x-axis is shown only up to 50 pN as all the simulations showed 0% beads bound before the rupture force reached 50 pN. Either 1 (left column), or 2 (right column) CBM constructs are attached to each bead. The 3 constructs per bead plots is shown in Figure 5-B in the main manuscript. Note that the difference between single and tandem profiles becomes more pronounced upon increasing the number of constructs/bead and with increased loading rate (see highlighted columns in Supplemental Table S5).*

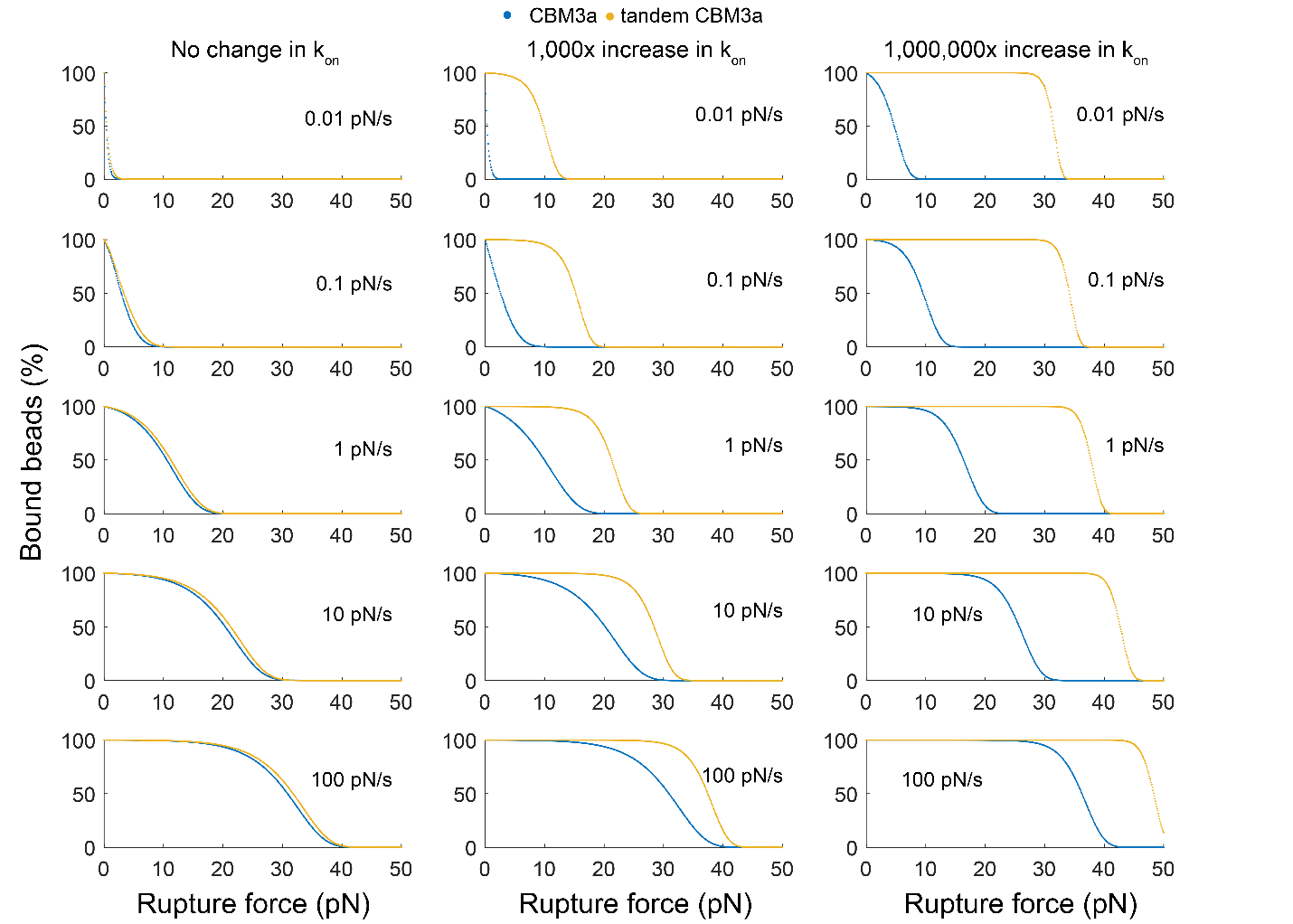

**Supplemental Figure S8:** Simulated percentage of CBM3a-functionalized beads bound to cellulose I polymers as a function of the applied rupture force (pN). Beads are functionalized in the simulations with either single (blue curves) or tandem (yellow curves) CBM constructs. A “no-rebind” WLC polymer model (see Supplemental Methods) was used in these runs where beads that become fully detached from cellulose are not allowed to rebind. Each row gives plots generated for a particular loading rate (from top to bottom: 0.01 pN/s, 0.1 pN/s, 1 pN/s, 10 pN/s, and 100 pN/s). The rupture force in the simulations went up to 200 pN; for visualization, the x-axis is shown only up to 50 pN as all the simulations showed 0% beads bound before the rupture force reached 50 pN (except the 10^6^ increase in k_on_ at 100 pN/s). Either no change in k_on_ (left column), a 1,000x increase in k_on_ (middle column), or a 10^6^ x increase in k_on_ (right column) was used in the simulations. Each bead had 3 constructs attached.

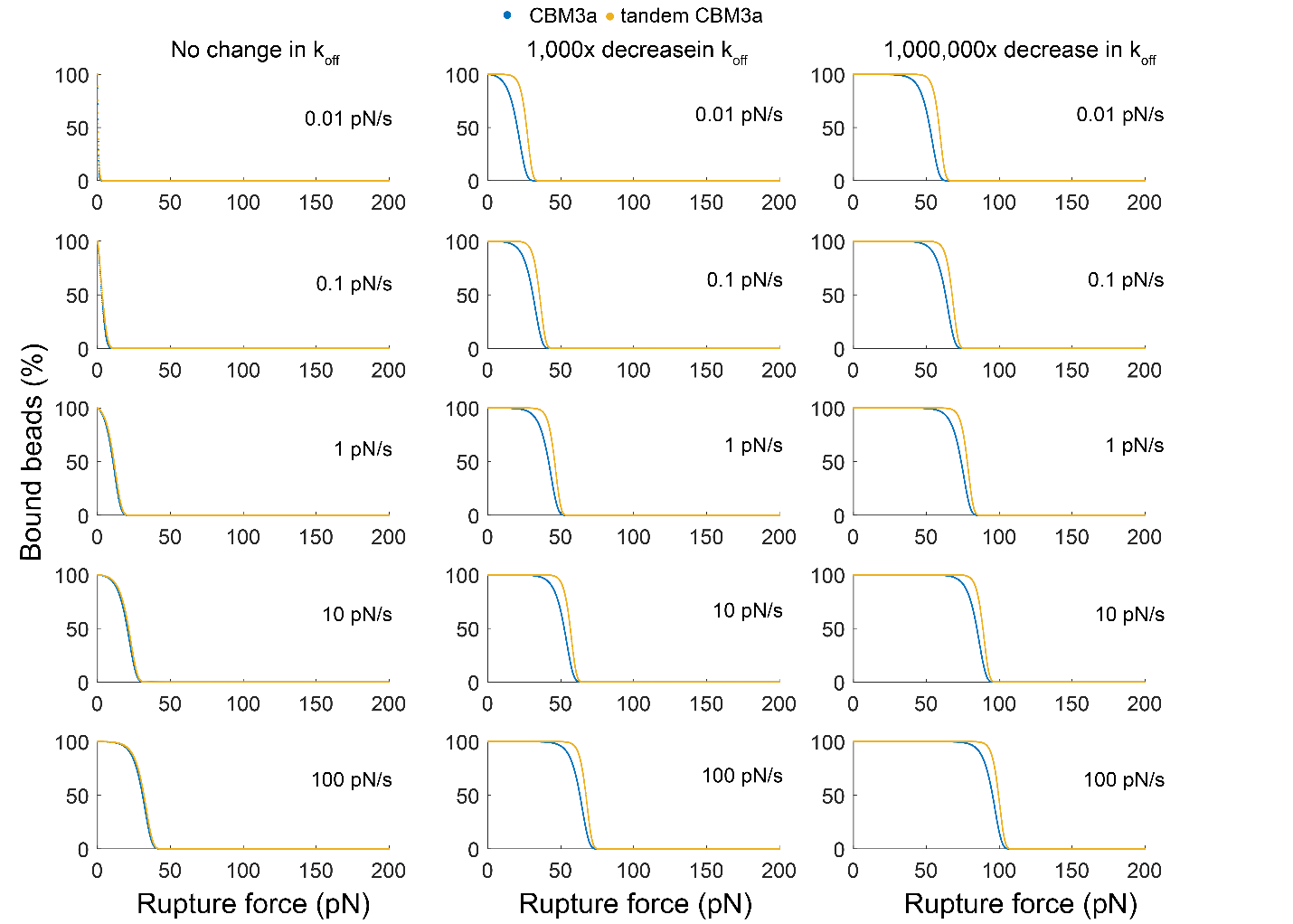

**Supplemental Figure S9**: Simulated percentage of CBM3a-functionalized beads bound to cellulose I polymers as a function of the applied rupture force (pN). Beads are functionalized in the simulations with either single (blue curves) or tandem (yellow curves) CBM constructs. A “no-rebind” WLC polymer model (see Supplemental Methods) was used in these runs where beads that become fully detached from cellulose are not allowed to rebind. Each row gives plots generated for a particular loading rate (from top to bottom: 0.01 pN/s, 0.1 pN/s, 1 pN/s, 10 pN/s, and 100 pN/s). The rupture force in the simulations went up to 200 pN. Either no change in k_off_ (left column), a 1,000x decrease in k_off_ (middle column), or a 1,000,000x decrease in k_off_ (right column) was used in the simulations. Each bead had 3 constructs attached.

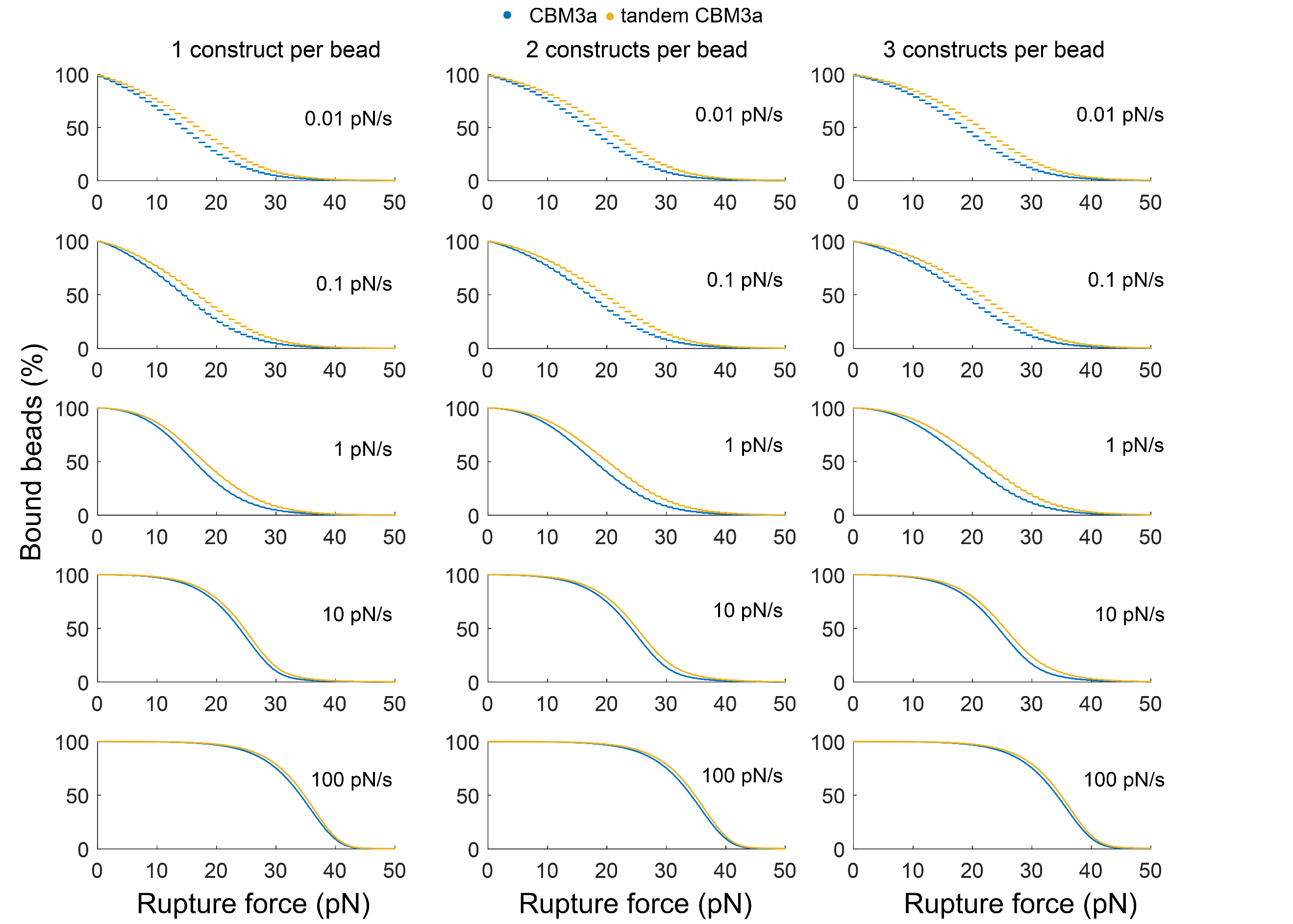

***Supplemental Figure S10****: Simulated percentage of CBM3a-functionalized beads bound to cellulose I polymers as a function of the applied rupture force (pN). Beads are functionalized in the simulations with either single (blue curves) or tandem (yellow curves) CBM constructs. A “can-rebind” WLC polymer model (see Supplemental Methods) was used in these runs where beads that become fully detached from cellulose are allowed to rebind. Each row gives plots generated for a particular loading rate (from top to bottom: 0.01 pN/s, 0.1 pN/s, 1 pN/s, 10 pN/s, and 100 pN/s). The rupture force in the simulations went up to 200 pN; for visualization, the x-axis is shown only up to 50 pN as all the simulations showed 0% beads bound before the rupture force reached 50 pN. Either 1 (left column), 2 (middle column), or 3 (right column) constructs are attached to each bead. Note that the difference between single and tandem profiles becomes less pronounced with increased loading rate (see orange columns in Supplemental Table S6).*

**Supplemental Table S1:** Nucleotide sequence of tandem His-Avi-GFP-CBM3a. The letters in bold black indicate the sequence for the His_8_ tag, followed by the TEV cleavage site (orange), Avi tag (blue), GFP domain (green), first linker domain (grey), first CBM3a domain (red), second linker domain (grey) and second CBM3a domain (red).

ATGGGA**CATCACCATCATCACCATCACCATGCATCCGAAAACCTGTACTTCCAGGGACTAAATGATATATTTGAAGCTCAAAAAATCGAGTGGCACGAA**GCGATCGCC**TCCAAAGGTGAAGAACTGTTCACCGGTGTTGTTCCGATCCTGGTTGAACTGGACGGTGACGTTAACGGTCACAAATTCTCCGTTTCCGGTGAAGGTGAAGGTGACGCTACCTACGGTAAACTGACCCTGAAATTCATCTGCACCACCGGTAAACTGCCGGTTCCGTGGCCGACCCTGGTTACCACCCTGACCTACGGTGTTCAGTGCTTCTCCCGTTACCCGGACCACATGAAACAGCACGACTTCTTCAAATCCGCTATGCCGGAAGGTTACGTTCAGGAACGTACCATCTCCTTCAAAGACGACGGTAACTACAAAACCCGTGCTGAAGTTAAATTCGAAGGTGACACCCTGGTTAACCGTATCGAACTGAAAGGTATCGACTTCAAAGAAGACGGTAACATCCTGGGTCACAAACTGGAATACAACTACAACTCCCACAACGTTTACATCACCGCTGACAAACAGAAAAACGGTATCAAAGCTAACTTCAAAATCCGTCACAACATCGAAGACGGTTCCGTTCAGCTGGCTGACCACTACCAGCAGAACACCCCGATCGGTGACGGTCCGGTTCTGCTGCCGGACAACCACTACCTGTCCACCCAGTCCGCTCTGTCCAAAGACCCGAACGAAAAACGTGACCACATGGTTCTGCTGGAATTCGTTACCGCTGCTGGTATCACCCACGGTATGGACGAACTGTACAAA**GGTTTAAAC**GCGACTCCCACTAAAGGTGCCACTCCTACCAATACGGCGACTCCGACTAAGTCGGCAACGGCAACGCCCACTCGCCCCAGCGTACCGACCAATACTCCGACTAATACCCCGGCGAACACC**CTTAAG**GTAAGCGGTAACCTGAAGGTTGAATTTTATAACTCCAACCCAAGCGACACAACGAATAGCATCAATCCGCAGTTCAAAGTCACGAACACTGGCAGTTCAGCTATCGATCTGTCGAAACTGACCCTTCGTTACTACTATACGGTTGATGGCCAAAAAGATCAGACCTTTTGGTGCGACCATGCAGCAATCATCGGTAGCAATGGTTCTTATAACGGCATTACTTCTAATGTAAAAGGCACCTTTGTGAAGATGTCAAGTAGCACCAACAATGCTGATACCTACCTGGAAATTAGCTTCACGGGTGGCACACTTGAACCAGGAGCCCACGTCCAGATCCAGGGCCGTTTTGCGAAAAACGATTGGAGCAACTATACGCAATCAAACGATTATAGTTTCAAAAGCGCGTCTCAATTCGTAGAATGGGATCAGGTGACCGCATATTTGAACGGAGTGCTGGTTTGGGGGAAAGAACCAGGA**GGTTTAAAC**GCGACTCCCACTAAAGGTGCCACTCCTACCAATACGGCGACTCCGACTAAGTCGGCAACGGCAACGCCCACTCGCCCCAGCGTACCGACCAATACTCCGACTAATACCCCGGCGAACACC**CTTAAG**GTAAGCGGTAACCTGAAGGTTGAATTTTATAACTCCAACCCAAGCGACACAACGAATAGCATCAATCCGCAGTTCAAAGTCACGAACACTGGCAGTTCAGCTATCGATCTGTCGAAACTGACCCTTCGTTACTACTATACGGTTGATGGCCAAAAAGATCAGACCTTTTGGTGCGACCATGCAGCAATCATCGGTAGCAATGGTTCTTATAACGGCATTACTTCTAATGTAAAAGGCACCTTTGTGAAGATGTCAAGTAGCACCAACAATGCTGATACCTACCTGGAAATTAGCTTCACGGGTGGCACACTTGAACCAGGAGCCCACGTCCAGATCCAGGGCCGTTTTGCGAAAAACGATTGGAGCAACTATACGCAATCAAACGATTATAGTTTCAAAAGCGCGTCTCAATTCGTAGAATGGGATCAGGTGACCGCATATTTGAACGGAGTGCTGGTTTGGGGGAAAGAACCAGGA**TAA

**Supplemental Table S2:** Primer sequences for the insertion of the Avi tag between TEV cleavage site and GFP of pEC-GFP-CBM plasmids and removal of the histidine tag from the MBP-BirA plasmid.

| **Primer** | **Sequence** |
| --- | --- |
| Avi-insert_FW | GGACTAAATGATATATTTGAAGCTCAAAAAATCGAGTGGCACGAAGCGATCGCCTCCAAA |
| Avi-insert_BW | TTCGTGCCACTCGATTTTTTGAGCTTCAAATATATCATTTAGTCCCTGGAAGTACAGGTT |
| His-removal-BirA_FW | ACCATAGCATATGAAAATCGAAGAAGG |
| His-removal-BirA_BW | CGATTTTCATATGCTATGGTCCTTGTTG |

**Supplemental Table S3:** Molar extinction coefficients and molecular weights of the 4 prepared GFP-CBM constructs.

| **Construct** | **Molar extinction coefficient**  **(M^-1^cm^-1^)** | **Molecular weight**  **(kDa)** |
| --- | --- | --- |
| GFP-CBM3a | 64,415 | 52.865 |
| GFP-CBM64 | 76,905 | 45.753 |
| Tandem GFP-CBM3a | 99,950 | 74.80 |
| Tandem GFP-CBM64 | 124,805 | 60.584 |

**Supplemental Table S4:** Calculated effective K_D_’s for the binding of tandem CBM3a and CBM64 domains to cellulose I and PASC using a “corrected” WLC polymer model of multivalency (see main text for a description of the corrective factor incorporated here). K_D_ values are provided for several lengths of the disordered linker between the tandem CBM domains.

|  | | **45-aa linker (current)** | | **10-aa linker** | | **100-aa linker** | |
| --- | --- | --- | --- | --- | --- | --- | --- |
| **System** | **K_D,single_ (μM)** | **K_D,tandem_ (nM)** | **binding enhancement** | **K_D,tandem_ (nM)** | **binding enhancement** | **K_D,tandem_ (nM)** | **binding enhancement** |
| CBM3a + cellulose I | 0.28 | 0.56 | 502x | 0.12 | 2308x | 1.24 | 225x |
| CBM3a + PASC | 0.30 | 0.64 | 469x | 0.14 | 2154x | 1.43 | 210x |
| CBM64 + cellulose I | 0.29 | 0.60 | 485x | 0.13 | 2229x | 1.33 | 218x |
| CBM64 + PASC | 2.12 | 32.0 | 66x | 7.0 | 305x | 71.0 | 30x |

**Supplemental Table S5:** Calculated f_50_ or the rupture force (pN) at which 50% of CBM3a-functionalized beads remain bound to cellulose 1. A “no-rebind” WLC polymer model (see Supplemental Methods) was used in these runs where beads that become fully detached from cellulose are not allowed to rebind. Values are shown for different loading rates and different number of CBM constructs/bead. **^a^**	Difference between tandem CBMs f_50_ and single CBM f_50_.

|  | **1 construct/bead** | | | **2 constructs/bead** | | | **3 constructs/bead** | | |
| --- | --- | --- | --- | --- | --- | --- | --- | --- | --- |
| **Loading rate (pN/s)** | **single CBM f_50_ (pN)** | **tandem CBMs f_50_ (pN)** | **diff f_50_ (pN) ^a^** | **single CBM f_50_ (pN)** | **tandem CBMs f_50_ (pN)** | **diff f_50_ (pN) ^a^** | **single CBM f_50_ (pN)** | **tandem CBMs f_50_ (pN)** | **diff f_50_ (pN) ^a^** |
| 0.01 | 0.8 | 0.9 | 0.1 | 0.7 | 0.8 | 0.1 | 0.4 | 0.5 | 0.1 |
| 0.1 | 5.1 | 5.3 | 0.2 | 4.8 | 5.1 | 0.3 | 2.9 | 3.3 | 0.4 |
| 1 | 13.7 | 14.2 | 0.5 | 13.6 | 14.1 | 0.5 | 10.6 | 11.2 | 0.6 |
| 10 | 23.6 | 24.2 | 0.6 | 23.6 | 24.2 | 0.6 | 20.4 | 21.3 | 0.9 |
| 100 | 34.1 | 34.7 | 0.6 | 34.1 | 34.7 | 0.6 | 30.9 | 31.8 | 0.9 |

**Supplemental Table S6:** Calculated f_50_ or the rupture force (pN) at which 50% of CBM3a-functionalized beads remain bound to cellulose 1. A “can-rebind” WLC polymer model (see Supplemental Methods) was used in these runs where beads that become fully detached from cellulose are allowed to rebind. Values are shown for different loading rates and different number of CBM constructs/bead. **^a^**	Difference between tandem CBMs f_50_ and single CBM f_50_.

|  | **1 construct/bead** | | | **2 constructs/bead** | | | **3 constructs/bead** | | |
| --- | --- | --- | --- | --- | --- | --- | --- | --- | --- |
| **Loading rate (pN/s)** | **single CBM f_50_ (pN)** | **tandem CBMs f_50_ (pN)** | **diff f_50_ (pN) ^a^** | **single CBM f_50_ (pN)** | **tandem CBMs f_50_ (pN)** | **diff f_50_ (pN) ^a^** | **single CBM f_50_ (pN)** | **tandem CBMs f_50_ (pN)** | **diff f_50_ (pN) ^a^** |
| 0.01 | 14.1 | 16.3 | 2.2 | 16.2 | 19.1 | 2.9 | 18.1 | 20.6 | 2.5 |
| 0.1 | 14.3 | 16.9 | 2.6 | 16.9 | 19.9 | 3.0 | 18.3 | 21.1 | 2.8 |
| 1 | 16.3 | 18.0 | 1.7 | 17.9 | 20.1 | 2.2 | 19.2 | 21.5 | 2.3 |
| 10 | 23.8 | 24.6 | 0.8 | 24.1 | 25.1 | 1.0 | 24.3 | 25.4 | 1.1 |
| 100 | 34.2 | 34.8 | 0.6 | 34.2 | 34.8 | 0.6 | 34.2 | 34.8 | 0.6 |

#### Supplemental Information References

1.	A. Sethi, B. Goldstein, S. Gnanakaran, Quantifying intramolecular binding in multivalent interactions: A Structure-Based synergistic study on Grb2-Sos1 complex. *PLoS Comput. Biol.* **7**, 1–13 (2011).

2.	T. Travers, *et al.*, Combinatorial diversity of Syk recruitment driven by its multivalent engagement with FcεRIγ. *Mol. Biol. Cell* **30**, 2331–2347 (2019).

3.	N. Rawat, P. Biswas, Size, shape, and flexibility of proteins and DNA. *J. Chem. Phys.* **131** (2009).

4.	H. X. Zhou, Loops in proteins can be modeled as worm-like chains. *J. Phys. Chem. B* **105**, 6763–6766 (2001).

5.	V. S. Chauhan, S. K. Chakrabarti, Use of nanotechnology for high performance cellulosic and papermaking products. *Cellul. Chem. Technol.* **46**, 389–400 (2012).

6.	J. R. Faeder, M. L. Blinov, W. S. Hlavacek, “Rule-Based Modeling of Biochemical Systems with BioNetGen” in *Systems Biology*, I. V Maly, Ed. (Humana Press, 2009), pp. 113–167.

7.	L. A. Harris, *et al.*, BioNetGen 2.2: Advances in rule-based modeling. *Bioinformatics* **32**, 3366–3368 (2016).

8.	M. Hackl, *et al.*, Acoustic force spectroscopy reveals subtle differences in cellulose unbinding behavior of carbohydrate-binding modules. *Proc. Natl. Acad. Sci.* **119**, e2117467119 (2022).

9.	D. Jayachandran, *et al.*, Engineering and characterization of carbohydrate‐binding modules for imaging cellulose fibrils biosynthesis in plant protoplasts. *Biotechnol. Bioeng.* **120**, 2253–2268 (2023).

10.	G. I. Bell, Models for the specific adhesion of cells to cells. *Science (80-. ).* **200**, 618–627 (1978).
